# A multiscale functional map of somatic mutations in cancer integrating protein structure and network topology

**DOI:** 10.1101/2023.03.06.531441

**Authors:** Yingying Zhang, Alden K. Leung, Jin Joo Kang, Yu Sun, Guanxi Wu, Le Li, Jiayang Sun, Lily Cheng, Tian Qiu, Junke Zhang, Shayne Wierbowski, Shagun Gupta, James Booth, Haiyuan Yu

**Author notes:** These authors contributed equally to this work (Yingying Zhang, Alden K. Leung).

## Abstract

A major goal of cancer biology is to understand the mechanisms underlying tumorigenesis driven by somatically acquired mutations. Two distinct types of computational methodologies have emerged: one focuses on analyzing clustering of mutations within protein sequences and 3D structures, while the other characterizes mutations by leveraging the topology of protein-protein interaction network. Their insights are largely non-overlapping, offering complementary strengths. Here, we established a unified, end-to-end 3D structurally-informed protein interaction network propagation framework, NetFlow3D, that systematically maps the multiscale mechanistic effects of somatic mutations in cancer. The establishment of NetFlow3D hinges upon the Human Protein Structurome, a comprehensive repository we compiled that incorporates the 3D structures of every single protein as well as the binding interfaces of all known protein interactions in humans. NetFlow3D leverages the Structurome to integrate information across atomic, residue, protein and network levels: It conducts 3D clustering of mutations across atomic and residue levels on protein structures to identify potential driver mutations. It then anisotropically propagates their impacts across the protein interaction network, with propagation guided by the specific 3D structural interfaces involved, to identify significantly interconnected network “modules”, thereby uncovering key biological processes underlying disease etiology. Applied to 1,038,899 somatic protein-altering mutations in 9,946 TCGA tumors across 33 cancer types, NetFlow3D identified 1,4444 significant 3D clusters throughout the Human Protein Structurome, of which ~55% would not have been found if using only experimentally-determined structures. It then identified 26 significantly interconnected modules that encompass ~8-fold more proteins than applying standard network analyses. NetFlow3D and our pan-cancer results can be accessed from http://netflow3d.yulab.org.

## Main

Somatically acquired mutations are one of the major sources driving tumorigenesis^1^. Computational approaches have been developed to assign pathogenicity scores to given mutations, indicating their phenotypic effects on an organism^2–8^. Complementary to these approaches, understanding the mechanisms driven by each mutation—from altering genomic sequences to changing key amino acid residues to dysregulating relevant cellular pathways—is key to developing effective therapeutic strategies. Efforts have been made to interpret the effects of mutations at specific scales^9–17^. Some studies focus on the molecular effects and look for spatial clustering of mutations within critical regions of proteins^9–15,18,19^. Others focus on cancer pathways and look for significantly mutated subnetworks of proteins^16,17,20^. Studies at the molecular and pathway levels offer complementary insights into the underlying mechanisms of cancer.

At the 3D protein structural level, the spatial clustering of mutations on 3D protein structures can reveal functionally important protein regions and can thus assist in identifying cancer driver mutations^9–15,19^. Given that the overwhelming majority of somatic mutations in cancer are non-functional passengers^21^, 3D clustering analysis narrows down potential driver mutations and thus significantly boost the signal-to-noise ratio. However, previous 3D clustering algorithms either limit their scope to the experimentally-determined structures^9,11,12,15^, or specifically focus on single proteins^10–12,14^ or protein-protein interaction (PPI) interfaces^22,23^. No approach yet fully examines the 3D structures of every single protein as well as the binding interfaces of all known PPIs in humans, leaving many spatial clusters yet to be identified. The bottleneck has been the limited coverage of 3D structural information: only ~36% of single proteins and ~6% of known PPIs in humans have experimentally-determined structures^24^. Nonetheless, recent breakthroughs in deep learning technologies for highly accurate 3D structure prediction, covering both single proteins^25–29^ and multi-protein complexes^30–34^, are rapidly filling these gaps.

At the PPI network level, various methods have been developed to identify significantly mutated subnetworks by integrating genetic mutation data with network topology^16,17,35,36^. These strategies have revealed many key pathways and protein complexes in cancer. Furthermore, sophisticated analyses can construct a hierarchy of altered subnetworks^20,37,38^, offering a nuanced, multi-layered perspective on the cancer-related biological processes across various subnetwork levels.

The insights gained from 3D protein structural level and PPI network level methodologies are largely non-overlapping, thereby offering complementary strengths. Integrating these methodologies is key to comprehensively delineate cancer mechanisms. Here, we compiled the Human Protein Structurome–a comprehensive repository of 3D structural data that covers every single protein as well as all known PPIs in humans–by leveraging recent deep-learning-based protein structural information prediction strategies^24,25^, upon which we established NetFlow3D, a unified framework that integrates methodologies across 3D structural and PPI network levels to systematically map the multiscale functional effects of somatic mutations across atomic, residue, protein and network scales.

NetFlow3D integrates 3D structural information and PPI network topology in a comprehensive, end-to-end process: (i) Initially, it identifies potential driver mutations through 3D clustering analysis on protein structures, and only propagates these 3D clustering signals across the PPI network, significantly improving our signal-to-noise. (ii) NetFlow3D recognizes that proteins often interact with different partners using distinct 3D structural interfaces. It, therefore, uses this information to weight the impact of spatially clustered mutations at a specific PPI interface on different interaction partners differently (anisotropic) (Supplementary Fig. 1).

NetFlow3D is one coherent and unified framework that integrates information across all four (atomic, residue, protein, and network) levels, thereby reinforcing the confidence of discoveries at all levels: for example, 3D clustering of mutations across atomic and residue levels allows network propagation of only likely driver mutations and pinpoints their specific impacts on different interaction partners; while network propagation and topological analysis further boost confidence in those 3D mutation clusters that are significantly interconnected within the same module, and shed light on complex biological processes underlying disease etiology. This end-to-end 3D-structurally-informed network propagation framework allows us to identify a much greater number of likely functional mutations and a more extensive range and larger scale of disease-associated network modules, which demonstrates molecular and clinical significance but cannot be identified by traditional methods.

By applying NetFlow3D to a TCGA pan-cancer dataset across 33 cancer types, we generated a multiscale functional map of somatic mutations in cancer: (i) Our analysis revealed over twice as many significant 3D clusters throughout the Human Protein Structurome as those identified using only experimentally-determined structures. (ii) NetFlow3D identified 26 significantly interconnected modules within the PPI network, each characterized by densely interconnected 3D clusters. These modules incorporate ~8 times more proteins than those identified through standard PPI network analyses that do not incorporate any 3D protein structural information. To enhance the accessibility of the NetFlow3D tool and our pan-cancer study results for the broader biomedical community, we built a user-friendly interactive web server (http://netflow3d.yulab.org/) where researchers can directly apply NetFlow3D to their own datasets and interactively browse our TCGA pan-cancer results.

## Results

### NetFlow3D maps the functional effects of somatic mutations across multiple scales

We compiled and processed a TCGA pan-cancer dataset of 1,038,899 somatic protein-altering mutations across 9,946 tumor samples spanning 33 cancer types (Fig. 1a; Methods). Of these mutations, 82% were expected to change only one or a few amino acid residues in the encoded proteins (i.e. missense mutations and in-frame indels), and are thus collectively referred to as in-frame mutations. Without further biological contexts, it is particularly difficult to interpret the varying downstream functional effects based on these subtle changes to the protein sequences.

**Fig. 1.**
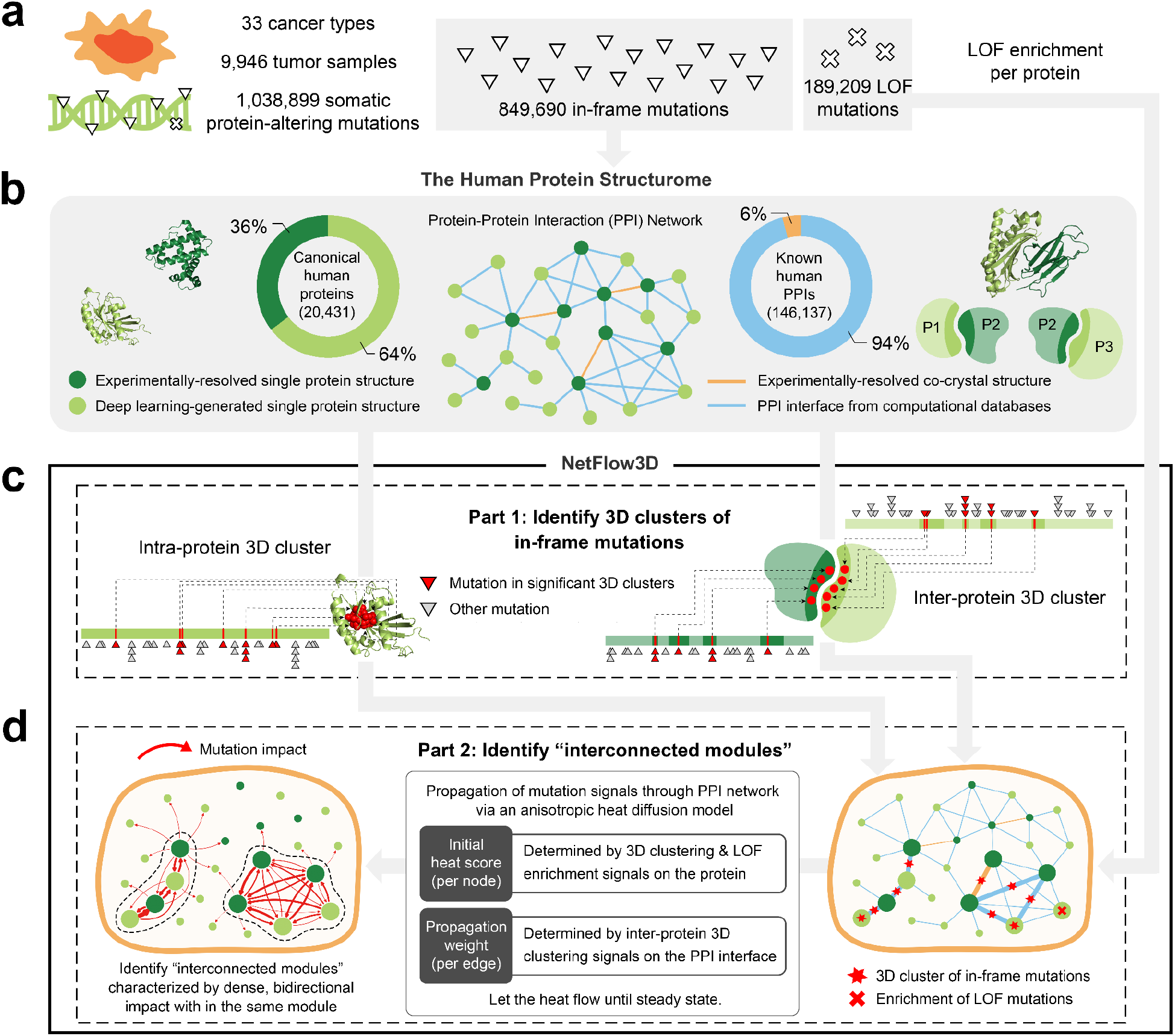
Framework of mapping the multiscale functional effects of somatic mutations. (**a**) Preprocessed TCGA pan-cancer mutation dataset. (**b**) Overview of the Human Protein Structurome. For 3D structures of single proteins, in addition to the 20,431 canonical isoforms illustrated, the Human Protein Structurome also incorporates 165,328 non-canonical isoforms. (**c**) The first part of NetFlow3D, a 3D clustering algorithm for identifying both intra- and inter-protein 3D clusters of in-frame mutations. (**d**) The second part of NetFlow3D, a network propagation model for interpreting the pathway-scale effects of mutations in 3D clusters by identifying interconnected modules.

Mounting evidence has demonstrated the efficaciousness of identifying functional in-frame mutations by detecting spatial clusters on 3D protein structures^9–15,18,19^. In order to achieve full-coverage spatial mapping of mutations on 3D protein structures, we compiled a comprehensive repository that contains the structures of all human proteins as well as the binding interfaces of all known human PPIs and available multi-protein complex structures, which we named “the Human Protein Structurome” (Fig. 1b; Methods). Importantly, the 3D structural data of 64% of canonical human proteins and 94% of known human PPIs were generated by recent deep-learning approaches, including AlphaFold2^25^ and PIONEER^24^, which were not available to previous 3D clustering algorithms.

The first part of NetFlow3D is a 3D clustering algorithm that identifies spatial clusters of in-frame mutations throughout the entire Human Protein Structurome (Fig. 1c; Methods). Our algorithm looks for both 3D clusters within single proteins (intra-protein 3D clusters) and 3D clusters spanning interacting proteins (inter-protein 3D clusters). Unlike most existing 3D clustering algorithms, (i) our method models the varying local background mutation rate across the genome by accounting for replication timing, mRNA expression level, HiC chromatin compartment, local GC content, and local gene density, an approach adapted from MutSigCV^39^ (Methods). This differs from the common practice in many 3D clustering algorithms that determine the significance of 3D clusters by randomly shuffling mutations within the same protein structure. (ii) Our method determines the physical contact between every pair of amino acid residues by accounting for their varying 3D distances across all available structures instead of solely based on a single snapshot represented by one structure (Methods).

The second part of NetFlow3D employs a heat diffusion model adapted from HotNet2^16^ to propagate 3D clustering signals (“heat”) through the PPI network (“diffusion”) (Fig. 1d; Methods). Importantly, our method goes beyond traditional PPI network analyses by incorporating 3D structural information in two crucial aspects: (i) NetFlow3D assigns an initial heat score to each node (protein) based on the 3D clustering signals on that protein, unlike traditional PPI network analyses that rely on gene mutation frequency or other gene-level statistics, thereby significantly boosting the signal-to-noise ratio. (ii) When NetFlow3D propagates heat from one node to its neighbors (representing the impact of 3D mutation clusters), it assigns additional propagation weight to the edges (PPIs) that have 3D mutation clusters on their corresponding PPI interfaces. This strategy is grounded in the “edgetic effect” of functional missense mutations, indicating that mutations at the interface are more likely to disrupt the corresponding PPI than non-interface mutations. This effect has been observed in both germline^40–42^ and somatic mutations (Supplementary Fig. 2; Supplementary Table 1). NetFlow3D’s weighted propagation strategy differs from traditional PPI network analyses that typically treat all edges connected to a given node as equal. Subsequently, NetFlow3D identifies “interconnected modules” within the network, i.e., subnetworks characterized by densely interconnected 3D clusters. To be in the same module, two proteins, *u* and *v*, should both have substantial 3D clustering signals that significantly impact each other. This method is designed to prevent the formation of “star graphs”, which are centered around well-studied cancer proteins but include surrounding proteins with minimal 3D clustering signals and biological relevance.

As a complement to the first and second parts that focus on in-frame mutations, NetFlow3D also accounts for loss-of-function (LOF) mutations. These mutations, which drastically alter protein sequences, are generally less specific about where they occur within protein structures to exert their effect. Therefore, NetFlow3D evaluates the enrichment of LOF mutations scattered across the entire sequence of each protein, and incorporates these protein-specific LOF enrichment signals as additional initial heat scores into the heat diffusion model in the second part (Fig. 1a and 1d; Methods).

Overall, NetFlow3D maps the functional effects of somatic mutations across multiple scales: from (i) atomic-resolution 3D clustering of mutations, to (ii) identification of protein residues as functionally significant by significant 3D clusters, to (iii) the perturbation of key proteins/PPIs, to (iv) the dysregulation of network modules and cellular pathways. This multiscale functional map provides rich insights for understanding cancer mechanisms driven by the subtle genomic mutations in somatic tissues.

### Significant intra- and inter-protein 3D clusters throughout the Human Protein Structurome

We applied the 3D clustering algorithm in NetFlow3D to the 849,690 somatic in-frame mutations in the TCGA pan-cancer dataset. This analysis led to the identification of 7,634 significant intra-protein 3D clusters and 6,810 significant inter-protein 3D clusters throughout the Human Protein Structurome (Fig. 2a; Supplementary Table 2). Notably, 60% of intra-protein clusters and 50% of inter-protein clusters were identified using 3D structural data from deep learning databases. For example, within the 3D structure of PPP2R5B protein generated by AlphaFold 2, we identified an intra-protein 3D cluster composed exclusively of rarely mutated residues (i.e., mutated in no more than two tumor samples) (Fig. 2b). These residues would not have been identified through individual analysis. Impressively, 99.1% of residues in our significant 3D clusters do not exhibit significant recurrent mutations when analyzed individually (Supplementary Fig. 3a). However, these infrequently-mutated residues demonstrate a significant enrichment for catalytic residues (Supplementary Fig. 3b). The use of AlphaFold 2-generated structures was crucial in identifying these potentially functional, yet infrequently mutated residues in proteins without experimentally-resolved structures. Moreover, single protein structures alone (even if covering every human protein) are still not enough for the comprehensive identification of all 3D clusters. This is because many driver mutations accumulate at the binding interfaces of cancer-related PPIs^22,45,48^. Only looking at individual proteins will split inter-protein 3D clusters into smaller fragments on individual proteins, making them harder to identify. This is demonstrated by the fact that, among the identified residues within our significant inter-protein 3D clusters, 55.8% would not have been identified if we only searched for significant intra-protein 3D clusters. Such situations are exemplified by an inter-protein cluster on the PPI interface between RHOC and ARHGAP1 proteins, as revealed by PIONEER (Fig. 2c). These results highlight the importance of knowing PPI interfaces, which are mostly generated by our deep learning framework PIONEER, in identifying potential driver mutations. Overall, 91.6% of TCGA tumor samples with somatic in-frame mutations have at least one mutation incorporated by our significant 3D clusters, demonstrating the thoroughness of our 3D cluster identification.

**Fig. 2.**
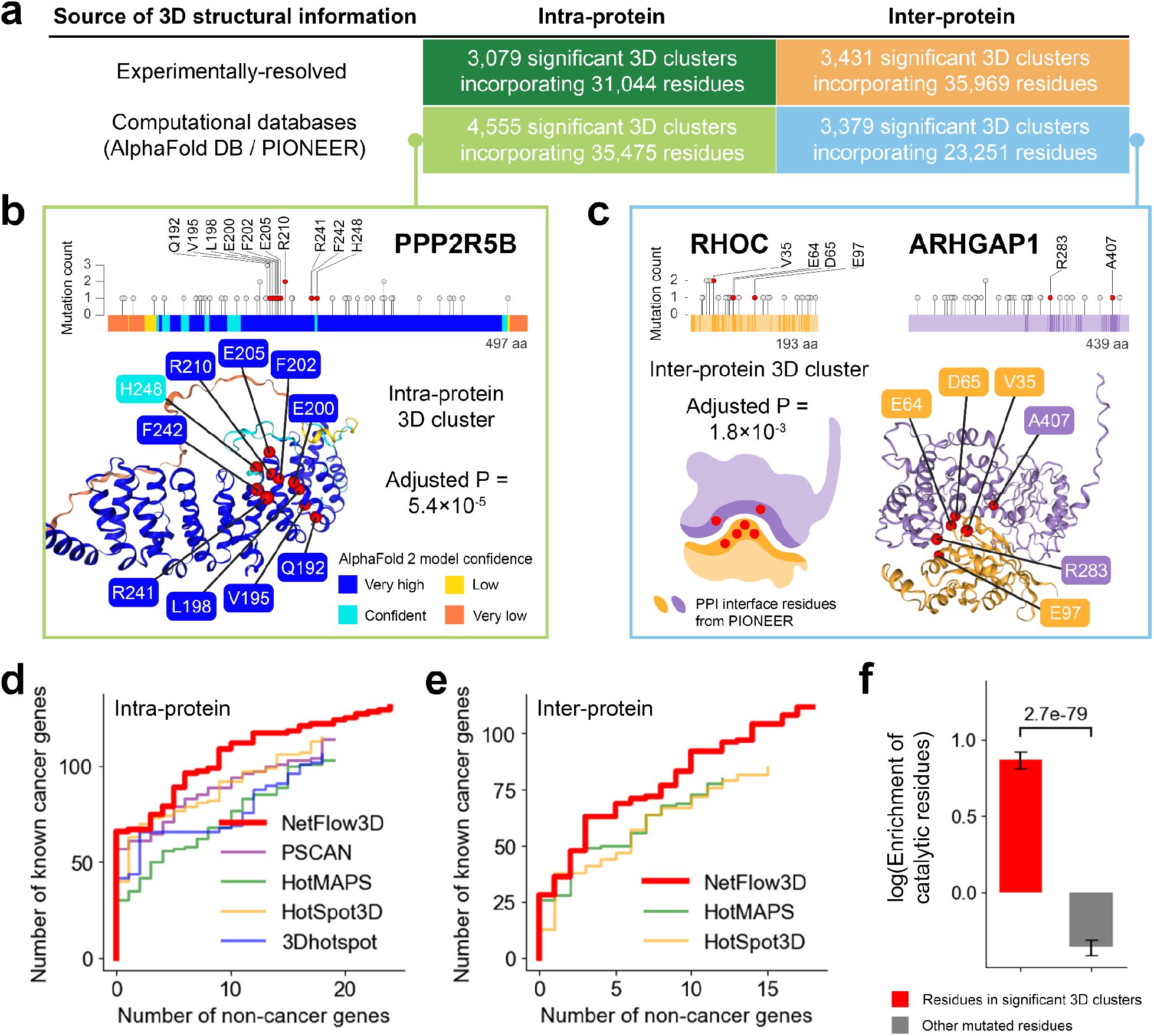
Description of 3D clusters identified by NetFlow3D and performance evaluation. (**a**) Summary of the significant intra- and inter-protein 3D clusters identified by NetFlow3D. (**b**) An example of a significant intra-protein 3D cluster identified on the AlphaFold 2-generated structure of PPP2R5B protein. All mutations incorporated by this cluster are on the residues with “very high” or “confident” model confidence. (**c**) An example of a significant inter-protein 3D cluster on the binding interface between RHOC and ARHGAP1 proteins. The PPI interface is resolved by PIONEER. For visualization purposes, a 3D structure of this protein complex is generated using AlphaFold Multimer to highlight the significant inter-protein 3D cluster. (**d**) Analysis of enrichment for catalytic residues in 3D clustering results from NetFlow3D. (**e-f**) Performance comparison between NetFlow3D and state-of-the-art 3D clustering algorithms. Performance curves are drawn for the top 1-500 genes, ranked by each algorithm based on the highest scoring 3D cluster on each gene.

We then evaluated the performance of NetFlow3D and compared it with four state-of-the-art 3D clustering algorithms^9–11,13^ which represent major sources of 3D cluster identification (Methods). We applied each algorithm to the same TCGA pan-cancer dataset, and compared the 3D clusters identified by different algorithms. Considering that (i) some algorithms only focus on intra-protein clusters^10,11^ while some others identify both^9,13^, and (ii) some algorithms only use experimentally-determined structures^9,11^ while some others also include comparative protein structure models^10,13^, we therefore make coherent comparisons by (i) assessing the intra- and inter-protein clusters separately, and (ii) limiting the comparisons to the 3D clusters identified on experimentally-resolved structures. Genes were ranked by each algorithm according to the highest score obtained from all the 3D clusters present on them. As a result, within the same number of top genes ranked by each algorithm, NetFlow3D-ranked genes consistently include a higher number of known cancer genes listed by the Cancer Gene Census (CGC)^49,50^ (Supplementary Table 3) as well as a lower number of non-cancer-associated genes^51–53^ (Supplementary Table 4), demonstrating our advanced sensitivity and specificity (Fig. 2d-e). This was further validated using an independent pan-cancer dataset from the Catalogue of Somatic Mutations in Cancer (COSMIC)^50^, where NetFlow3D maintained its leading performance (Supplementary Fig. 4; Methods).

Beyond 3D clustering algorithms, we benchmarked NetFlow3D against other methods for identifying cancer driver mutations, including single-residue-based (“hotspot”) and whole-gene-based methods. The test unit size of 3D clustering algorithms falls between these two extremes. Notably, NetFlow3D outperforms these methods, demonstrating the highest precision and recall (Supplementary Fig. 5). While the hotspot method is highly precise, it lacks power when background mutation rates are low or sample sizes are small. The whole-gene-based method, which considers the entire gene as the test unit, can dilute statistical power and lacks precision when only specific regions within the gene are responsible for driving cancer. In contrast, our 3D clustering algorithm in NetFlow3D provides flexible test unit sizes at submolecular resolution, achieving a balance of higher precision and better power.

Overall, the 3D clusters identified by NetFlow3D demonstrate a significant enrichment for catalytic residues^43^ (fold enrichment: 2.4, p-value: 3.7e-48, two-sided Z-test), while mutated residues outside these clusters exhibit a significant depletion (fold enrichment: 0.7, p-value: 4.7e-13, two-sided Z-test) (Fig. 2f; Methods). This pattern remains robust and is not sensitive to variations in p-value cutoffs (Supplementary Fig. 6). Notably, this robust pattern is consistent across 3D clusters identified from both experimentally-determined structures and deep-learning-generated 3D structural data (Supplementary Fig. 7a). Considering the intrinsic bias of inter-protein clusters towards functional residues, as PPI interface residues are known to be enriched for such residues^22–24,44–47^, we specifically excluded these inter-protein clusters from our analysis and strictly focused on intra-protein clusters. Our refined analysis shows that the previously identified pattern persists (Supplementary Fig. 7b). Moreover, proteins involved in our 3D clusters demonstrate a significant enrichment for known cancer genes (fold enrichment: 1.7, p-value: 3.6e-18, two-sided Z-test), whereas proteins not involved in any 3D clusters show a significantly depletion (fold enrichment: 0.56, p-value: 2.9e-14, two-sided Z-test) (Fig. 3a). This pattern is robust, remaining consistent across a range of p-value cutoffs (Supplementary Fig. 8).

**Fig. 3.**
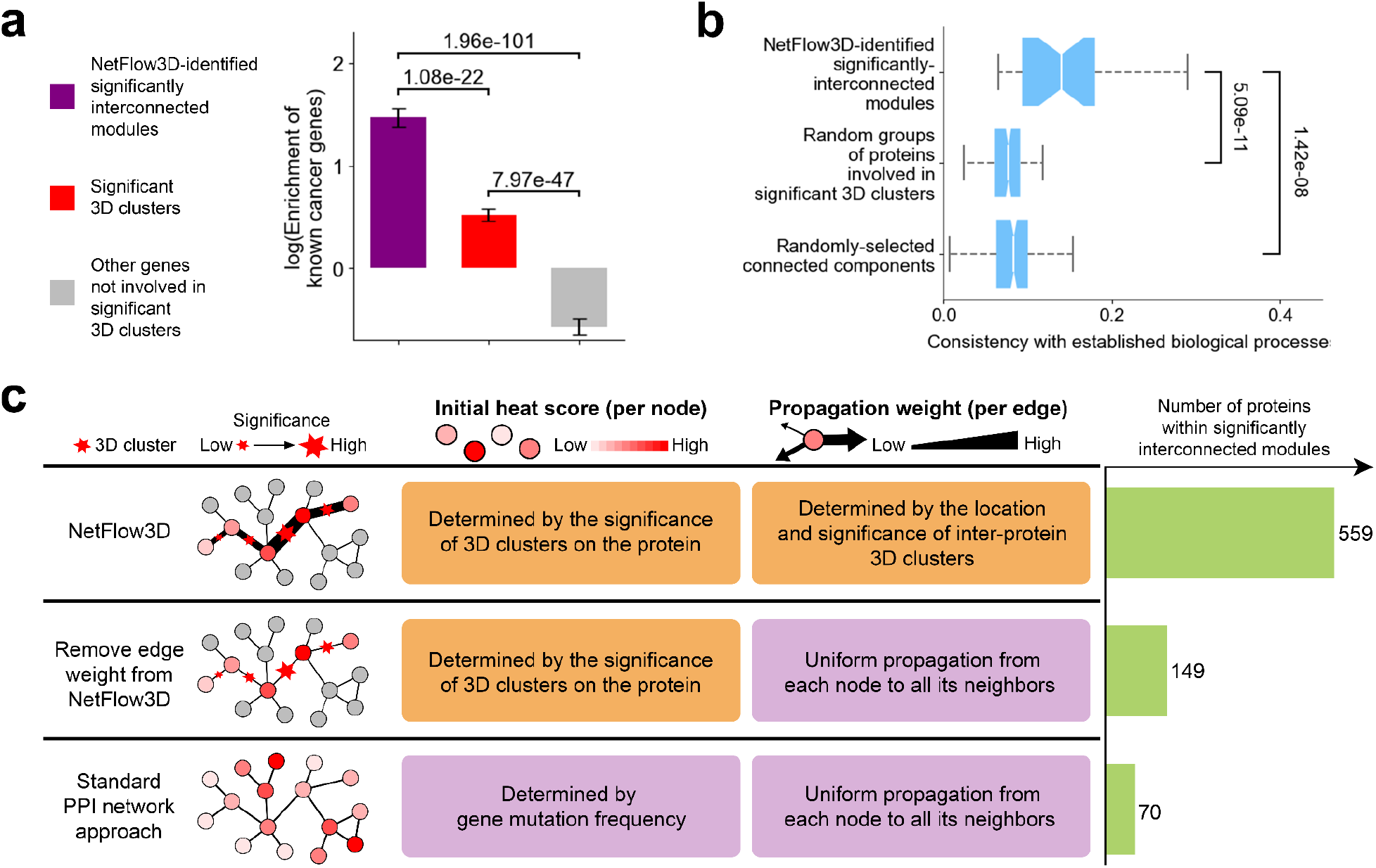
The benefits of integrating 3D structural information and PPI Network topology end-to-end over using either alone. (**a**) Enrichment analysis for known cancer genes conducted on three sets of proteins supported by different levels of evidence. The p-values were calculated using two-proportion Z-tests. (**b**) Consistency between NetFlow3D-identified significantly interconnected modules and established biological processes, compared to random background. The p-values for comparisons are provided by the Mann-Whitney U test. (**c**) Results from systematically removing two key strategies that NetFlow3D used to incorporate 3D structural information via nodes and edges.

### Importance of our end-to-end integration of 3D structural information and PPI network topology

The critical innovation of NetFlow3D over previous methods lies in its seamless, end-to-end integration of 3D structural information with PPI network topology. To underscore the unique insights this integration provides, we compared the outcomes of NetFlow3D with those from methods that use only information in either 3D protein structures or PPI network topology.

The advantage of our end-to-end integration over solely relying on 3D protein structural information manifests in two key aspects. First, the dense interconnections among 3D clusters within the same module further reinforce their validity, bolstering confidence in molecular-level discoveries. This is evidenced by the observation that proteins within NetFlow3D-identified significantly interconnected modules (Supplementary Table 5) exhibit a 2.6-fold higher enrichment for known cancer genes compared to those identified solely by significant 3D clusters, even though the latter already demonstrate significant enrichment (Fig. 3a). Second, by extending the analysis beyond identifying crucial 3D structural regions within proteins, the propagation of 3D mutation clustering signals throughout the PPI network provides deeper insights into the dysregulated biological processes underlying tumorigenesis. This is demonstrated by the observation that significantly interconnected modules identified by NetFlow3D align more closely with established biological processes^54–58^ than do random groups of those proteins with significant 3D clusters which were organized to match the NetFlow3D-identified modules in number and sizes (Fig. 3b). However, this closer alignment is not just an outcome of the PPI network’s topology, as randomly selected connected components with matched number and sizes show significantly lower consistency with established biological processes (Fig. 3b). Thus, it’s the effective integration of molecular-level 3D clustering information and the PPI network’s topology that plays a key role in uncovering critical biological processes that are potentially central to cancer development.

The advantage of our end-to-end integration over the methods relying solely on PPI network topology is the significant improvement in statistical power. This improvement is demonstrated by the outcomes of systematically removing the two key strategies that NetFlow3D used to incorporate 3D structural information via nodes and edges (Fig. 3c; Methods). Initially, the edge weight in NetFlow3D, determined by 3D clustering signals on PPI interfaces, was removed, leading to uniform propagation from each node to all its neighbors. As a result, the significantly interconnected modules identified thereafter contain ~¼ of the proteins found in the original NetFlow3D-identified significantly interconnected modules. Next, the initial heat scores assigned to each node, determined by the 3D clustering signals on each protein, was replaced by gene mutation frequency. This further change fully reverted the original NetFlow3D framework to a standard PPI network approach. Consequently, the significantly interconnected modules identified thereafter contain only ~⅛ of the proteins identified by the original NetFlow3D framework.

### Biological significance of NetFlow3D-identified significantly interconnected modules

We benchmarked NetFlow3D-identified significantly interconnected modules (hereafter called “NetFlow3D modules”) against well-established cancer signaling pathways^59^ (positive controls) (Supplementary Table 6) and Gene Ontology (GO) biological processes (BPs)^54–58^ (background reference) (Supplementary Table 7). Enrichment analysis for known cancer genes demonstrated that NetFlow3D modules exhibit enrichment levels comparable to those of well-established cancer pathways and significantly surpass those found in GO BPs (Fig. 4a). Furthermore, we analyzed mutation patterns within each entity—whether a NetFlow3D module, a well-known cancer pathway, or a GO BP—by calculating enrichment for two distinct mutation categories: (i) mutations within significant 3D clusters, and (ii) all mutations (Methods). Consequently, well-known cancer pathways and NetFlow3D modules consistently demonstrate pronounced enrichment trends for both mutation categories, with a particularly striking increase when switching from all mutations to the mutations within significant 3D clusters (Fig. 4b). In contrast, GO BPs exhibit no obvious trend of enrichment for all mutations and a much compromised enrichment for those within significant 3D clusters, with only a minor increase when contrasting the two mutation categories. Notably, upon splitting NetFlow3D modules into two groups based on whether they contain known cancer genes, the mutation patterns across the two groups are strikingly consistent (Fig. 4c), both resembling well-known cancer pathways (Fig. 4b). In contrast, GO BPs present a different picture: even those GO BPs that include known cancer genes exhibit much weaker mutation enrichment trends for both mutation categories. Meanwhile, GO BPs lacking known cancer genes display virtually no trend of mutation enrichment at all (Fig. 4c).

**Fig. 4.**
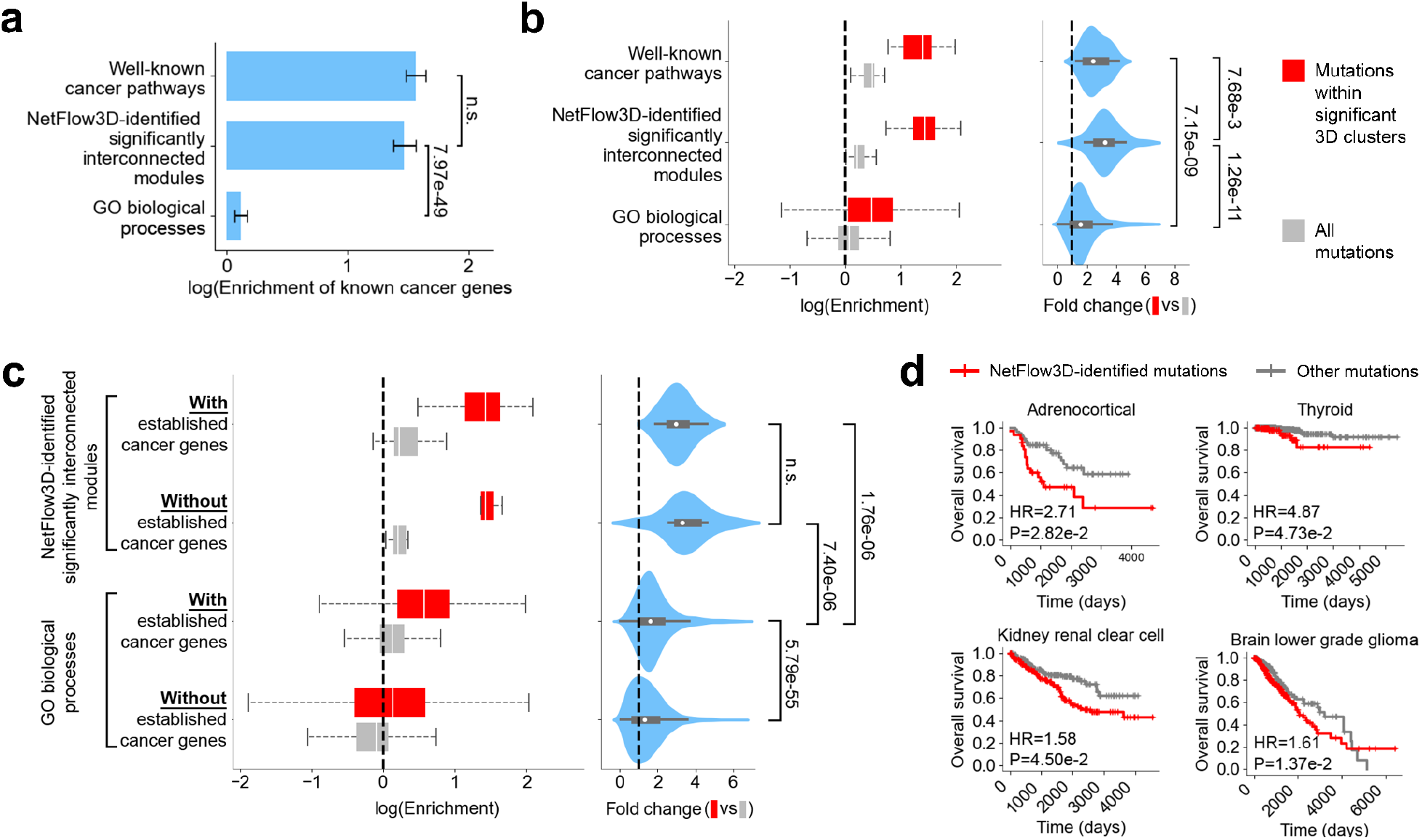
Evaluating the biological significance of NetFlow3D-identified significantly interconnected modules. (**a**) Enrichment comparison for known cancer genes among well-known cancer pathways, NetFlow3D modules, and GO biological processes, with p-values derived from two-proportion Z-tests. (**b**) Examination of mutation patterns within each well-known cancer pathway, NetFlow3D module, and GO biological process. The p-values for comparisons were obtained using the Mann-Whitney U test. (**c**) Comparison of mutation patterns between entities with versus without known cancer genes. This comparison was only applicable to NetFlow3D modules and GO biological processes. (**d**) Association of NetFlow3D-identified potential driver mutations with patients’ survival. HR stands for hazard ratio.

To demonstrate downstream molecular consequences of NetFlow3D-identified mutations, we evaluated their statistical association with protein abundance. Initially, we performed linear regression analysis, controlling for gene-specific and tissue-specific baseline expression levels, as well as clinical covariates including sex, age, tumor stage, and TMB (Supplementary Note). This analysis revealed significant associations between the presence of NetFlow3D-identified mutations and protein abundance (p-value: 1.6e-9). In contrast, no significant association was observed when conducting the same analysis using other mutations not identified by NetFlow3D (p-value: 0.70). For a more detailed perspective, we conducted a fine-grained analysis comparing protein abundance for each gene in each cancer type, under scenarios with and without NetFlow3D-identified mutations. As a control, we repeated the analysis using other mutations. Our results revealed a significantly higher proportion of cases with differential protein abundance for NetFlow3D-identified mutations than for other mutations (Supplementary Fig. 9).

To further evaluate the impact of genes with NetFlow3D-identified mutations on cellular fitness, we utilized core fitness (CF) genes identified from genome-scale CRISPR-Cas9 screens in 324 human cancer cell lines spanning 30 cancer types^60^. We analyzed the enrichment of these core fitness genes in NetFlow3D results. Our results consistently showed that, across various cancer types, genes with NetFlow3D-identified mutations are most enriched for core fitness genes (Supplementary Fig. 10). Additionally, genes with mutations identified in isolation by our 3D clustering analysis also showed significant enrichment in every cancer type, whereas genes without mutations in 3D clusters did not exhibit significant enrichment in any cancer type.

To demonstrate the clinical significance of NetFlow3D findings, we compared the overall survival between patients with somatic in-frame mutations in our preprocessed TCGA dataset, grouping them by whether their mutations were identified by NetFlow3D (Methods). We used a Cox regression model to evaluate the statistical association between NetFlow3D-identified mutations and patient survival, controlling for clinical covariates including age, sex, tumor stage, and tumor mutational burden (TMB). Our analysis revealed significant negative survival associations across multiple cancer types, including Thyroid carcinoma (THCA), Kidney renal clear cell carcinoma (KIRC), Adrenocortical carcinoma (ACC), and Brain Lower Grade Glioma (LGG) (Fig. 4d). The hazard ratios (HR) derived from the Cox model coefficients were consistently greater than 1.5 across all four cancer types (Supplementary Table 8).

Next, we assessed NetFlow3D’s capability to uncover novel insights beyond known cancer genes. Remarkably, 80% (447 out of 559) of the proteins identified within NetFlow3D modules are not encoded by known cancer genes listed in the CGC. Moreover, NetFlow3D-identified mutations in these non-CGC curated genes exhibited significant survival impacts in THCA and KIRC (Supplementary Fig. 11a; Methods). Furthermore, even after removing 3D clustering and LOF enrichment signals from known cancer genes and subsequently re-applying our 3D structurally-informed network propagation framework, the newly identified significantly interconnected modules still cover 23 out of the 26 original NetFlow3D modules (Supplementary Fig. 11b; Supplementary Note).

### A pan-cancer functional map of somatic mutations across scales

Applying NetFlow3D to the TCGA pan-cancer dataset has yielded a multiscale functional map of somatic mutations in cancer (Fig. 5). From a biological perspective, this map encompasses a broad spectrum of cellular processes and functions, spanning well-established cancer pathways, components that are increasingly-recognized through recent evidence, and biological entities with less-characterized roles in cancer (Supplementary Table 9). (i) Well-established cancer pathways. Examples include p53 signaling, regulation of apoptosis, regulation of E2F-dependent transcription, and intracellular signaling cascades like Ras, PI3K, mTOR, and TGF-β. (ii) Increasingly recognized pathways and protein complexes. Examples include Rho GTPase signal transduction^61^, chromatin remodeling (e.g. PRC2^62^, MLL complex^63^), immune processes (e.g. antigen processing and presentation^64^), and DNA repair mechanisms^65^ (e.g. interstrand cross-link repair and DNA non-homologous end joining). (iii) Biological entities with less-characterized roles in cancer. Examples include protein K11-linked ubiquitination^66^, eIF2 activity^67,68^, TRiC^69^, TFIID complex^70^, and calcineurin^71^. Notably, 46% of the NetFlow3D modules do not contain any known cancer genes listed in CGC. These modules are particularly intriguing as they represent new opportunities for cancer pathway exploration. Their significance has been underscored by the observation that these modules not only show mutation patterns strikingly similar to those of well-established cancer pathways (Fig. 4b-c) but also exhibit significant survival impacts (Supplementary Fig. 11a).

**Fig. 5.**
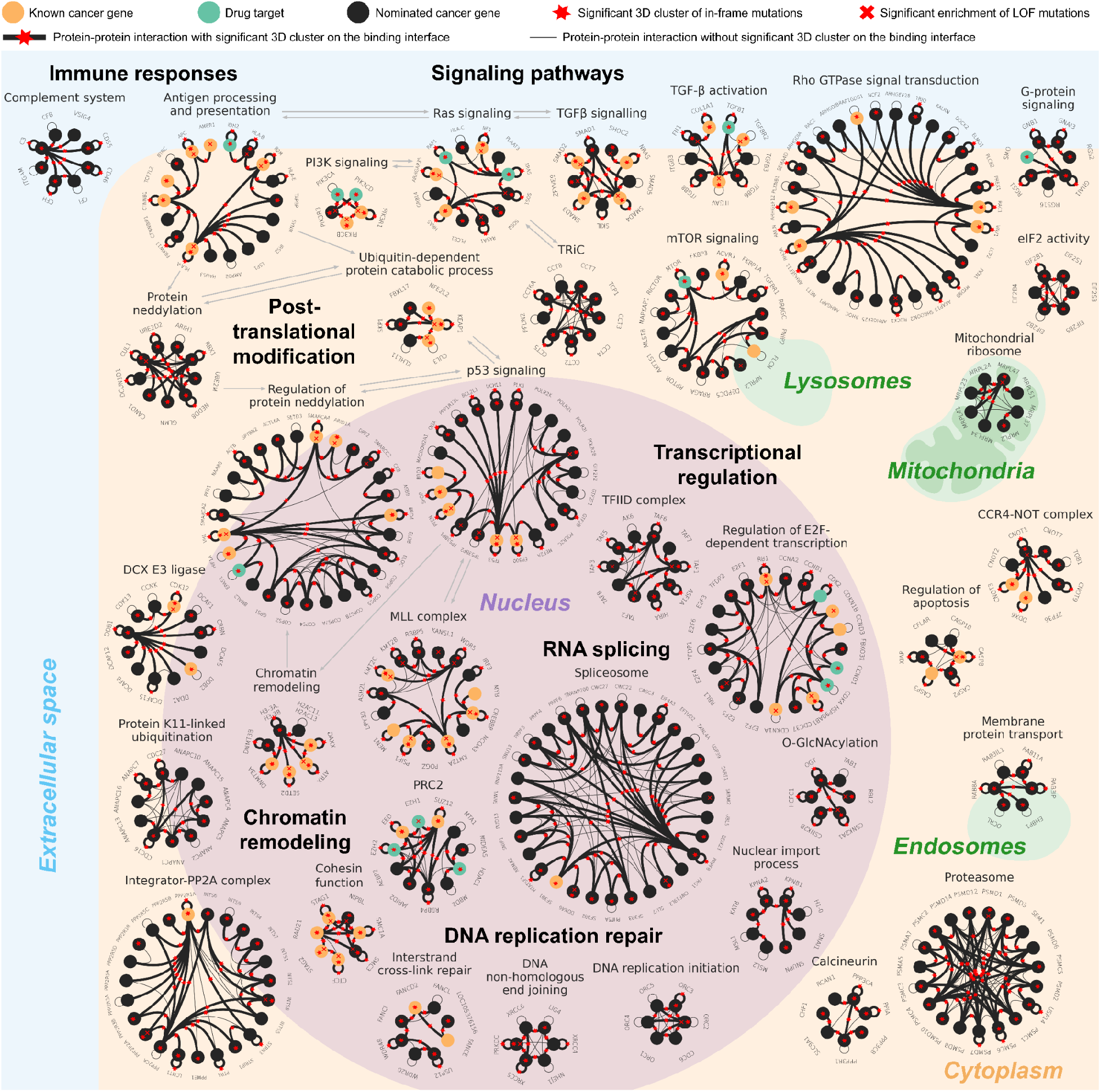
A multiscale functional map of somatic mutations in cancer. All the significantly interconnected modules identified by NetFlow3D were displayed here. The largest module is illustrated as the 11 core biological entities incorporated by this module, with direct mutation impacts between these entities represented by gray arrows. Importantly, red stars on the edges and the bolding of these edges indicate the presence of significant 3D mutation clusters on the binding interfaces between interacting protein pairs. Drug targets were labeled according to the full list of FDA-approved drugs and their corresponding targets provided in Supplementary Table 10.

Shifting the focus to the methodological advancements, this map generated by NetFlow3D not only aligns with key discoveries from traditional PPI network analyses, but also reveals novel insights achieved through integrating 3D structural information. Specifically, the map’s composition is threefold: (i) Components that are also identifiable via traditional PPI network approaches. For example, our map includes a vast majority of biological entities identified by HotNet2^16^, such as pathways like p53, PI3K, and KEAP1-NFE2L2. It also encompasses protein complexes such as MLL, cohesin, and SWI/SNF, as well as “linker genes” including regulators of Ras signaling and elements of MAPK signaling. Furthermore, our map also includes the complement system, as identified by Olcina et al.^72^ in their analysis of 69 cancer mutation datasets using HotNet2. (ii) Components emerging from combining PPI network topology and orthogonal data/analyses. For example, Wang et al.^73^ integrated PPI network topology with GWAS and identified the spliceosome. Gupta et al.^74^ integrated PPI network topology with gene co-expression network and external pathway annotations such as KEGG/Reactome/GO/IPA and identified Rho GTPase signal transduction. (iii) Components uniquely identified by NetFlow3D through integrating PPI topology with 3D structural information, such as PP2A, CCR4-NOT complex, mitochondrial ribosome, and calcineurin, etc. This distinct category highlights the novel insights provided by the end-to-end integration of the local spatial organization of mutations on 3D protein structures and their global topological organization in the network.

### Integrator-PP2A complex

To showcase how NetFlow3D reveals new insights into cancer biology, we presented one NetFlow3D module as an example, which corresponds to two established biological entities: the integrator complex^75^ and the PP2A complex^76^ (Fig. 6a). These two biological entities work collaboratively: PP2A is recruited to transcription sites by the integrator complex, where PP2A functionally counteracts CDKs-driven cell-cycle progression, thereby maintaining cell homeostasis^77–80^ (Fig. 6b).

**Fig. 6.**
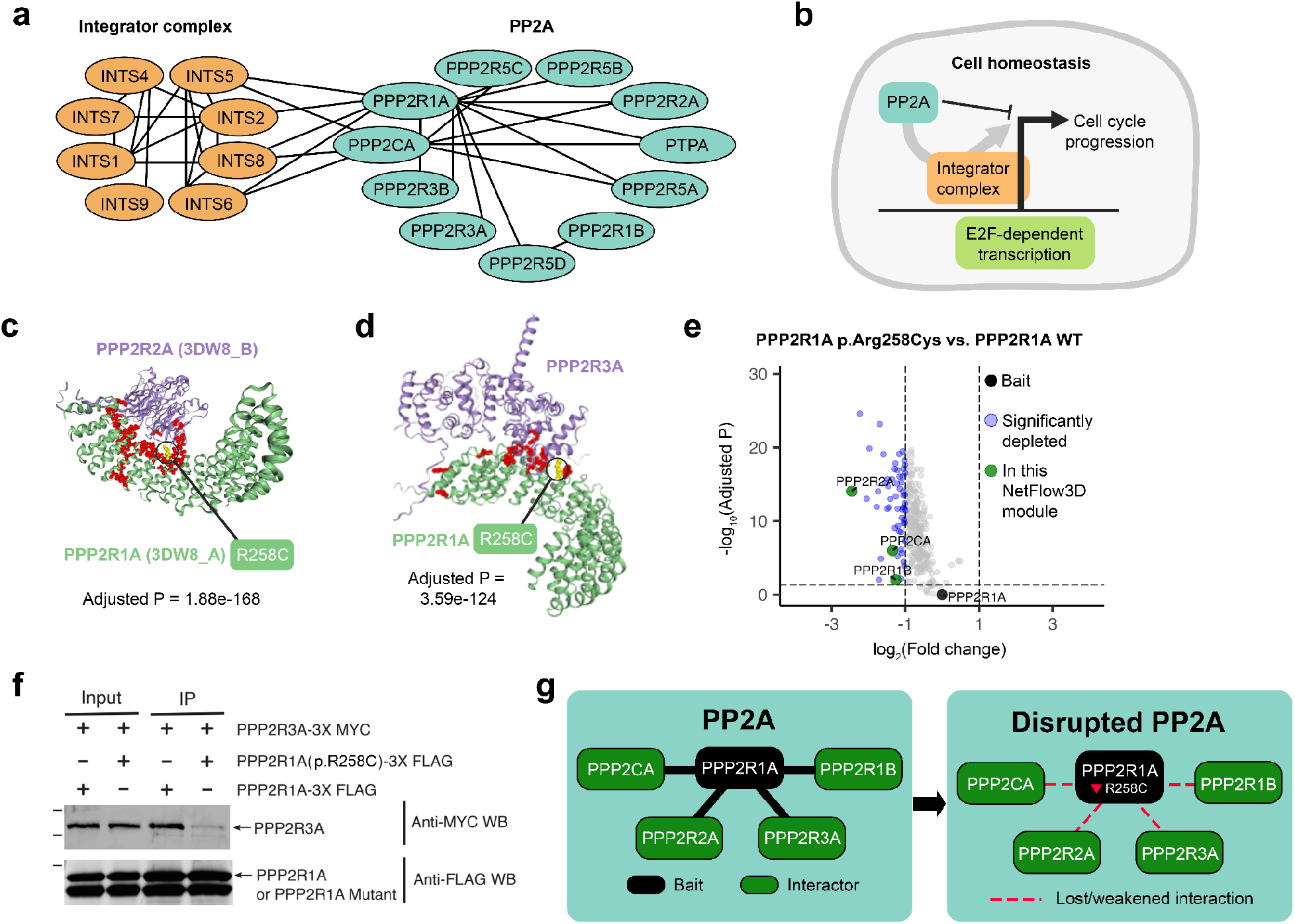
Illustration of a NetFlow3D module showcasing biological insights and experimental validation as proof-of-concept. (**a**) The biological entities incorporated by this module include the integrator complex and the PP2A complex. (**b**) Functionality of these entities: PP2A is recruited to transcription sites by the integrator complex, where PP2A functionally counteracts cell-cycle progression, thereby maintaining cell homeostasis. (**c-d**) A potential driver mutation highlighted in this module, PPP2R1A p.Arg258Cys, identified based on the significant 3D clusters at the binding interfaces with PPP2R2A and PPP2R3A. (**e**) TMT-IP-MS-based quantitative proteomics analysis confirming that PPP2R1A p.Arg258Cys mutation diminished PPP2R1A’s interactions with almost all of its interactors. The volcano plot summarizes the quantitative results for identified interactors that co-purify with PPP2R1A mutant (p.Arg258Cys) compared with PPP2R1A wildtype. Significantly depleted interactors were colored in blue, with fold change > 2, adjusted p-value < 0.05. PPP2R1A (bait) was colored in black, while other proteins from this module were colored in green, all of which are PP2A subunits. (**f**) Co-immunoprecipitation confirming that PPP2R1A p.Arg258Cys disrupted PPP2R1A–PPP2R3A interaction in HEK 293T cells. (**g**) Summary of experimentally validated disrupted PPP2R1A subnetwork.

Towards the identification of new candidate cancer driver mutations within this module, we focused on the p.Arg258Cys mutation in the PPP2R1A protein. NetFlow3D identified this mutation as being part of the significant 3D clusters at the binding interfaces between PPP2R1A/PPP2R2A proteins and PPP2R1A/PPP2R3A proteins (Fig. 6c-d). In our TCGA study, this mutation originated from a patient with uterine corpus endometrial carcinoma (UCEC). Additionally, mutations at the PPP2R1A codon 258 have been observed in serous and endometrioid carcinomas as reported in several non-TCGA studies^81–83^. Our quantitative-proteomics-based TMT-IP-MS and co-immunoprecipitation experiments showed that PPP2R1A p.Arg258Cys mutation diminished PPP2R1A’s interactions with almost all of its interactors (Fig. 6e). Particularly, it disrupted the interactions with other PP2A subunits within this module (Fig. 6e-g). Therefore, it is plausible to speculate that PPP2R1A p.Arg258Cys mutation diminished PP2A function by disrupting its subunit interactions. Given the previous evidence showing that the inactivation or inhibition of PP2A promotes cancer development^84–86^, our experimental validation of the disrupted PPP2R1A subnetwork (Fig. 6g) caused by the PPP2R1A p.Arg258Cys mutation underscores the value of NetFlow3D in identifying novel cancer driver mutations and illuminating potential tumorigenic mechanisms.

Importantly, NetFlow3D’s ability to identify the PP2A complex hinges upon our end-to-end integration of 3D structural information with PPI network topology. On one hand, PP2A was completely overlooked using the standard PPI network approach. On the other hand, >90% of PP2A subunits do not contain any single-residue hotspots, indicating that relying solely on mutation recurrence fails to capture this full biological entity. In contrast, by utilizing 3D structural insights from our Human Protein Structurome, NetFlow3D successfully identified significant 3D clusters on every PP2A subunit within this module, affirming their significant association with cancer individually. Furthermore, the dense interconnectivity among these significant 3D clusters, as revealed by NetFlow3D, further reinforces the overall functional significance of the PP2A complex in cancer biology.

## Discussion

Our work demonstrates the effective integration of 3D protein structural information with PPI network topology as achieved by NetFlow3D, our end-to-end 3D structurally-informed network propagation framework. This integration provides unique insights that can not be gained from each component in isolation. NetFlow3D applied 3D clustering analysis across the entire Human Protein Structurome, which not only identified >100-fold more potentially functional residues than using the single-residue-based hotspot method, but also discovered over twice as many significant 3D clusters compared to traditional 3D clustering analysis using only experimentally-resolved structures. Moreover, our strategy of 3D-structurally-informed network propagation led to the identification of a much higher number of significantly interconnected modules. These modules not only incorporated ~8 times more proteins than those identified by standard PPI network analyses, but also demonstrated a 2.6-fold greater enrichment of known cancer genes compared to solely leveraging 3D structural information, thereby unveiling many potentially novel aspects of cancer biology that were previously unrecognized.

In addition to pan-cancer studies, NetFlow3D is also applicable to studies focusing on specific cancer types. It enables users to not only input somatic mutation data, but transcriptome and interactome data tailored to a particular cancer tissue context. NetFlow3D then applies its 3D clustering algorithm to a subset of 3D structural data in the Human Protein Structurome, filtered based on the context-specific expression profile. Given that our Structurome contains the 3D structures of all human protein isoforms, it offers a great capacity to adapt to a variety of cellular contexts. Following this, NetFlow3D propagates 3D clustering signals through a context-specific PPI network. Considering the current limitations in interactome data, where most PPIs are mapped in generic contexts such as using yeast or HEK293 cell lines, NetFlow3D addresses this by filtering the general human PPI network with context-specific transcriptome data, thus focusing on the subnetwork of genes that are actually expressed. Looking ahead, as experimentally-determined cell-type-specific interactome data become available, we anticipate further improvement in NetFlow3D’s performance for these targeted applications.

Furthermore, the core principles of NetFlow3D are not confined to somatic mutations in cancer, but can be extended to understanding germline variants in various diseases. Recent studies have demonstrated that permutation-based 3D clustering analysis, when applied to neurodevelopmental disorders^87,88^ and the Human Gene Mutation Database (HGMD)^19^, can effectively identify rare disease-associated variants. Adapting NetFlow3D to utilize the latest genome-wide models of germline mutation rates at base pair resolution^89,90^ represents an advancement to these approaches. Additionally, NetFlow3D’s context-specific analyses are particularly well suited for studying diseases that manifest in specific tissues or cell types.

Despite these strengths, NetFlow3D’s performance is limited by the quality of available 3D structural data, especially those generated by advanced deep learning algorithms. Our Human Protein Structurome now contains atomic-resolution 3D structures for all individual human protein isoforms. However, for most PPIs, the Structurome is limited to interface residue data. Advanced deep learning algorithms, including various AlphaFold-based methods (such as AlphaFold-Multimer^32^, AF2Complex^31^, and others^30,33,34^) have begun producing atomic-resolution 3D structures for multi-protein complexes. Yet, these methods are currently capable of producing high-confidence models for only a very limited subset of PPIs. Therefore, updating the PPI interfaces in our existing Structurome with these atomic-resolution structural models is still a considerable challenge. Continued advancements in these techniques are expected to extend their coverage, and we foresee further enhancement in NetFlow3D’s performance as we integrate these evolving resources.

NetFlow3D also has limitations in fully accounting for all types of driver mutations. This includes in-frame mutations that, despite not clustering on 3D protein structures, are still functional in cancer. For example, mutations impacting protein stability often occur within the core of proteins, altering function without targeting specific residues. Similarly, mutations in intrinsically disordered regions (IDRs) can markedly disrupt overall protein flexibility. Moreover, copy number variations (CNVs), structural variants (SVs), and noncoding mutations – especially those affecting regulatory elements^91–97^, contribute to altering gene dosage or expression, thereby diversifying cancer mechanisms. Expanding NetFlow3D to integrate these mutation types would improve its ability to offer a more complete understanding of cancer biology, representing a crucial area for future development.

## Methods

### Data collection

#### The Cancer Genome Atlas (TCGA)

3.6M somatic mutations across 10,295 tumor samples and 33 cancer types were obtained from the standard MC3 analysis^98^. We included an additional 178 tumor samples in the current TCGA program (https://portal.gdc.cancer.gov/), not covered by the MC3 dataset but provided by Chang et al.^1^. RNA-seq data were obtained from Repository on the GDC data portal^99^ (https://portal.gdc.cancer.gov/).

#### The Catalogue of Somatic Mutations in Cancer (COSMIC)

We obtained coding point mutations from genome-wide screens (including whole exome sequencing) under genome assembly GRCh37, along with sample data and cancer classification information from COSMIC release v98^50^ (https://cancer.sanger.ac.uk). All TCGA tumor samples were excluded from this dataset to ensure independence. We further filtered the dataset to retain only primary tumor samples, i.e., retaining those labeled as “primary” in the “TUMOUR_SOURCE” column and excluding “cell-line”, “xenograft”, “organoid culture”, or “short-term culture” in the “SAMPLE_TYPE” column. To eliminate redundancy in COSMIC, we used the drop_duplicates() function in pandas, with key identifiers including “CHROMOSOME”, “GENOME_START”, “GENOMIC_WT_ALLELE”, “GENOMIC_MUT_ALLELE”, and “TUMOUR_ID”. The cancer classification information in this dataset was aligned with TCGA projects using subject matter expertise. Tumor samples that could not align with any TCGA projects were categorized as having an unknown cancer type.

### Data preprocessing

#### VEP annotation

A single canonical effect per mutation was reported using Variant Effect Predictor (VEP) version 107^100^, following the approach used by Chang et al.^1^. Additionally, to evaluate the consequence of accounting for proteoform diversity, we conducted analysis on the same TCGA mutation dataset, but mapping each mutation to all possible protein isoforms. Details on this are provided in the Supplementary Note. According to VEP annotations, we only retained protein-altering mutations, including LOF mutations (“Consequence” column: frameshift_variant, stop_gained, stop_lost, start_lost, splice_acceptor_variant, splice_donor_variant, splice_donor_5th_base_variant) and in-frame mutations (“Consequence” column: missense_variant, inframe_deletion, inframe_insertion).

#### Excluding germline variants

According to VEP annotations, we removed mutations with non-zero allele frequencies in gnomAD^101^ (“gnomADe_AF” column), which were identified as germline variants present in the general population. The consequences of applying this filter are detailed in the Supplementary Fig. 12.

#### Excluding mutations in unexpressed genes

We defined expressed genes of a specific cancer type as those with RNA expression levels ≥1 FPKM in ≥80% of tumor samples within that cancer type. We only retained the mutations in those expressed genes of their cancer types. For the tumor samples of unknown cancer types, we only retained their mutations in the genes that are expressed in ≥80% of TCGA cancer types. Following the approach by Leiserson et al.^16^, mutations in 18 well-known cancer genes (AR, CDH4, EGFR, EPHA3, ERBB4, FGFR2, FLT3, FOXA1, FOXA2, MECOM, MIR142, MSH4, PDGFRA, SOX1, SOX9, SOX17, TBX3, WT1) that have low transcript detection levels were exempted from the aforementioned RNA expression filter. The consequences of applying this filter are detailed in the Supplementary Fig. 13.

#### UniProt ID mapping

We obtained the ID mapping data from UniProt^102^, which incorporates the mapping between UniProt IDs and VEP-annotated Ensembl gene, transcript, and protein IDs. We mapped each mutation to UniProt entries, initially based on their annotated Ensembl protein IDs, then sequentially using Ensembl transcript and gene IDs if protein IDs are not available.

After data preprocessing, the TCGA dataset yielded 1,038,899 somatic protein-altering mutations across 9,946 tumor samples in 33 cancer types. The COSMIC dataset yielded 571,789 somatic protein-altering mutations across 12,352 tumor samples that were aligned to 27 TCGA cancer types.

### Construction of the Human Protein Structurome

#### Data collection

Experimentally-determined structures were obtained from the Protein Data Bank^103,104^ (PDB, http://www.rcsb.org/), specifically focusing on asymmetric units. Predicted 3D structures of all human protein isoforms were obtained from the AlphaFold Protein Structure Database^25,105^ (AlphaFold DB), encompassing both 20,431 canonical isoforms (Fig. 1b), and 165,328 non-canonical isoforms. Interface residue data for 146k known human PPIs were obtained from PIONEER^24^.

#### Processing of experimentally-determined structures

Residue-level mapping between UniProt and PDB entries were obtained from the Structure Integration with Function, Taxonomy and Sequences^106,107^ (SIFTS). Based on the PDB structures, we constructed two undirected graphs G_1_=(V_1_,E_1_) and G_2_=(V_2_,E_2_). G_1_ describes the physical contacts between residues in the same polypeptide chains, while G_2_ describes the physical contacts between residues in different polypeptide chains. V_1_ includes the UniProt residues covered by at least one PDB structure. E_1_ is the set of residue pairs in the same polypeptide chains whose minimal three-dimensional (3D) distances among all relevant PDB structures are no larger than 6Å. The 3D distance between two residues in a given PDB structure is defined as the euclidean distance between their closest atoms in that structure. E_2_ is the set of inter-chain residue pairs whose minimal 3D distances are no larger than 9Å. V_2_ is the set of residues involved in E_2_. G_1_ and G_2_ were added to the Human Protein Structurome.

#### Processing of 3D structural data from deep learning algorithms

Using the atomic-resolution 3D protein structures from AlphaFold DB, we constructed G_3_=(V_3_,E_3_) following the same procedures used for constructing G_1_. Residues with all levels of model confidence in these structures were taken into account. G_3_ was added to the Human Protein Structurome. For the interface residue data from PIONEER, we used “very high” confidence predictions. This dataset was also added to the Human Protein Structurome.

### NetFlow3D: Identifying 3D clusters of mutated residues

#### 3D cluster identification based on atomic-resolution 3D protein structures

(i) Intra-protein clusters: In-frame mutations were mapped to G_1_ and G_3_ respectively. Vertices affected by these mutations and the edges between them were extracted as subgraph g_1_ and g_3_. C_1_ and C_3_ are the sets of connected components in g_1_ and g_3_, which were considered intra-protein clusters identified based on PDB and AlphaFold DB structures, respectively. We removed the intra-protein clusters in C_3_ that have at least one residue overlapping with any 3D clusters in C_1_ to avoid reporting redundant 3D clusters. (ii) Inter-protein clusters: We obtained 119,526 high-quality binary PPIs for Homo Sapiens from HINT^108^ (http://hint.yulab.org/), a dataset released in August 2021. We restricted our focus to these PPIs (denoted as H) for the inter-protein 3D cluster identification. e_2_ is a subset of edges in G_2_ whose endpoints are both affected by in-frame mutations. For a PPI between protein A and B, g_1A_=(v_1A_,e_1A_) and g_1B_=(v_1B_,e_1B_) are subgraphs extracted from g_1_ incorporating the vertices and edges in A and B, respectively. e_2AB_ is a subset of e_2_ where each edge connects one vertex in v_1A_ and one vertex in v_1B_. In the merged graph g_2AB_=(v_1A_∪v_1B_,e_1A_∪e_1B_∪e_2AB_), C_2AB_ is the set of connected components having at least one edge in 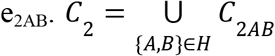 represents all inter-protein clusters identified based on PDB structures. Overall, C_structure_=C_1_∪C_2_∪C_3_ represents the set of 3D clusters identified based on atomic-resolution 3D protein structures.

#### 3D cluster identification based on PPI interface residue data from PIONEER

For a PPI between protein A and B, in-frame mutations were mapped to the interface residues, and the set of mutated interface residues was defined as an inter-protein 3D cluster, denoted as C_4AB_. *C*_*interface*_ = {*C*_4AB_ |{*A, B*} ∈ *H*} represents the set of inter-protein 3D clusters identified based on the PPI interface residue data from PIONEER.

### NetFlow3D: Background mutability model

To accurately model the background mutation rate (BMR) that varies extensively across the genome, we used a model that includes five covariates of mutation tendency: mRNA expression level, DNA replication timing, chromatin status as indicated by HiC mapping, local GC content, and gene density. The fundamental concept of this model originated from MutSigCV^39^: each gene *g* was positioned in a high-dimensional covariate space, estimating its local BMR based on its own silent and noncoding mutations, and, if necessary, those of its closest neighbors in this covariate space. Here, 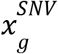 denotes the sum of silent and noncoding single nucleotide variants (SNVs) in gene g and its neighbors, and 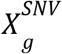 represents the total number of sequenced bases where silent and noncoding SNVs can occur in gene g and its neighbors. Consequently, the local BMR of coding SNVs in gene *g* is calculated as:

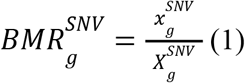

Similarly, 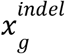 accounts for insertions and deletions (indels) within gene g and its neighbors, and 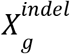 represents the total bases sequenced in the same regions. The local BMR for coding indels in gene g is calculated as:

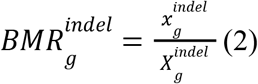

To estimate the expected number of in-frame and LOF mutations in gene *g*, we calculate the total number of covered bases in the coding region where mutation type *t* (missense, nonsense, splice site) can occur, denoted as 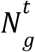. One base may contribute fractionally to multiple mutation types. For example, a covered C base might count 2/3 toward missense and 1/3 toward nonsense if mutations to A or G change the amino acid, while a mutation to T creates a stop codon. The probability of a random SNV in gene *g* falling into mutation type *t* is calculated as:

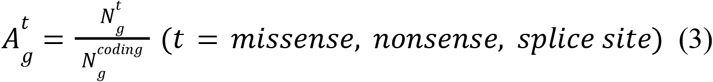

where 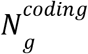 is the coding length of gene *g* in base pairs. Given that α = 9% (51,164 out of 56,031) of coding indels are in-frame and the rest are frameshift, we calculated the expected number of in-frame and LOF mutations in gene *g* as:

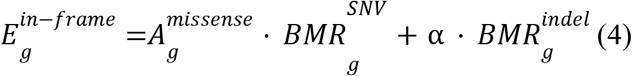

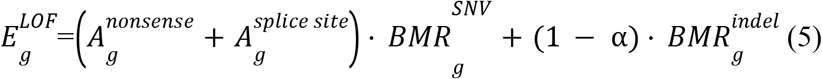

To avoid false positives due to exceedingly small local BMR in some genes, we set lower thresholds for 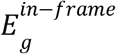 and 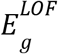 at the 0.01 quantile (1st percentile) of all 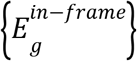 and all 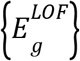, respectively. For a UniProt entry *u*, its expected number of in-frame and LOF mutations are calculated as:

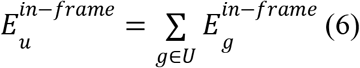

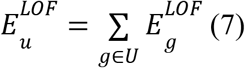

where *U* is the set of genes encoding this UniProt entry *u*. In cases where 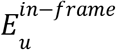 (or 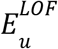) is absent, we adopted the median 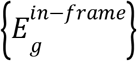 (or 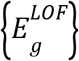) of all genes as default for that UniProt entry *u*.

### NetFlow3D: Determination of cluster significance

For an intra-protein 3D cluster *C* composed of *k* residues in UniProt entry *u*, the expected number of in-frame mutations across *n*_*p*_ patients in *C* is calculated as:

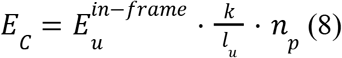

*l*_*u*_ is the length of UniProt entry *u* in amino acids.

For an inter-protein 3D cluster *C* spanning across the PPI interface of UniProt entry *u* and *v*, incorporating *k*_*u*_ residues in *u* and *k*_*v*_ residues in *v*, the expected number of in-frame mutations across *n*_*p*_ patients in *C* is calculated as:

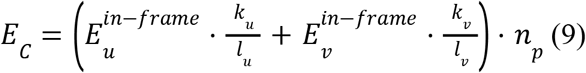

*O*_*C*_ denotes the observed number of in-frame mutations across *n*_*p*_ patients in *C*. The significance of *C* is determined by the one-sided p-value from Poisson test:

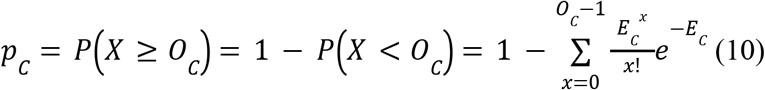

The Poisson test was applied to all 3D clusters. Bonferroni correction was separately applied to the 3D clusters in C_structure_ and C_interface_. 3D clusters with adjusted *p*_*C*_ <0.05 were considered significant. Additionally, we benchmarked these p-values via permutation tests (Supplementary Note), and observed a strong correlation between these p-values from NetFlow3D and those from permutation tests, with R^2^=0.75 (Supplementary Fig. 14).

### NetFlow3D: Protein-specific LOF enrichment signals

For a UniProt entry *u*, the expected number of LOF mutations across *n*_*p*_ patients is calculated as:

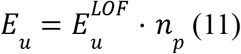

The significance of LOF enrichment in *u* is determined by the one-sided p-value from Poisson test:

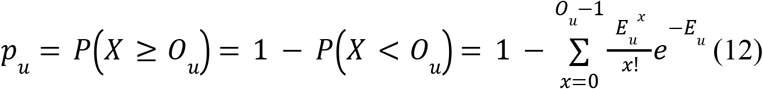

*O*_*u*_ denotes the observed number of LOF mutations across *n*_*p*_ patients in *u*. Bonferroni correction was applied, and the UniProt entries with adjusted *p*_*u*_ <0.05 were considered significantly enriched for LOF mutations.

### NetFlow3D: Network propagation model

#### Construction of the PPI network

The initial PPI network was built out of the aforementioned high-quality binary human PPIs from HINT. We filtered this network to encompass only genes expressed in any of the input cancer types. The aforementioned 18 well-known cancer genes with low transcript detection levels were considered expressed. The resulting PPI network was represented by an undirected graph G_PPI_=(V_PPI_,E_PPI_).

#### Heat definition

The initial amount of heat assigned to protein *u* was calculated as:

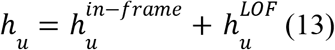

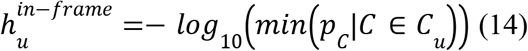

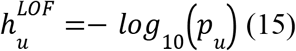

*C*_*u*_ denotes the set of 3D clusters that contain at least one residue in protein *u*. Both 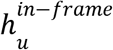 and 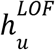 are constrained to a maximum value of 300 to prevent infinite heat scores that could have resulted from zero p-values. The initial heat distribution is described by a diagonal matrix *D*_*h*_ where the (*i, i*) entry is the amount of heat placed on protein *i*.

#### Heat transfer weight

At each time step, proteins in the PPI network pass to and receive heat from their neighbors, while retaining a fraction β of their heat. Notably, when a protein transfers its remaining 1-β fraction of heat to its neighbors, the heat is unevenly distributed. The amount of heat transferred along the edge between protein *i* and *j* is proportional to the weighting factor defined as:

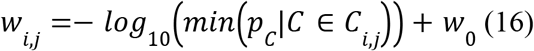

*C*_*i,j*_ denotes the set of inter-protein 3D clusters that are specific to the PPI between protein *i* and *j*. *w*_0_ = 1 is a baseline value, ensuring no edge has zero weight. *w*_*i,j*_ is also constrained to a maximum value of 300.

#### Heat diffusion and identification of interconnected modules

Once the initial heat assigned to each protein is determined, and the heat transfer weight along each edge is determined, the model is run until steady state is reached. If a unit of heat is placed on protein *j*, the net heat transferred from protein *j* to protein *i* is given by the (*i, j*) entry of the diffusion matrix *F* defined by:

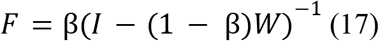

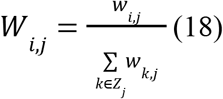

*Z*_*j*_ refers to the neighbors of protein *j*. The initial heat distribution is described by a diagonal matrix *D*_*h*_ where the (*i, i*) entry is the amount of heat placed on protein *i*. The exchanged heat matrix *E* is then defined by:

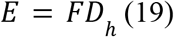

*E*(*i, j*) is the net heat transferred from protein *j* to protein *i*, given the initial heat *h*_*j*_ at protein *j*. We constructed a weighted directed graph based on E: If *E*(*i, j*) > δ, there was a directed edge from protein *j* to protein *i* of weight *E*(*i, j*). We then identified strongly connected components in this graph, which we term “interconnected modules”. A strongly connected component *C* in a directed graph is a set of vertices such that for every pair *u, v* of vertices in *C* there is a directed path from *u* to *v* and a directed path from *v* to *u*. Leiserson et al.^16^ have demonstrated that the identification of strongly connected components within a directed graph substantially reduced reporting “star graphs”, which are centered around well-studied, highly mutated cancer proteins, but include surrounding proteins with few mutations and little biological relevance. Our method strictly aligns with this principle, ensuring that the identified “interconnected modules” do not present with any one-way configurations. Thus, proteins in our “interconnected modules” are not merely passive recipients of influence from others’ 3D mutation clusters but also act as significant sources of influence.

#### Parameter determination

(i) Insulating parameter β: We used β = 0. 5, as used by HotNet2^16^. Edge weight parameter : The rationale behind selecting a δ is based on the fact that randomized data will typically not yield large “interconnected modules”. Therefore, choosing an appropriate value of δ can help identify “interconnected modules” that are statistically significant and likely to be biologically relevant. To generate a random undirected graph G_random_, we randomly swapped |E_PPI_| edge pairs in G_PPI_ while keeping the initial amount of heat on each protein fixed. The weighting factors {*w*_*i,j*_}were then randomly assigned to the newly swapped edges. Edge swapping was used to maintain the degree of each protein constant during the randomization process. We then applied the aforementioned heat diffusion model to G_random_ and identified the minimum δ such that all “interconnected modules” had size ≤ 5. We generated 20 such random directed graphs and identified a δ for each of them. We used the smallest value among these δ’s as the final value of δ. “Interconnected modules” exceeding a size of 5 were deemed significant, termed as “significantly interconnected modules” in our study.

### Implementing state-of-the-art 3D clustering algorithms

We applied four state-of-the-art 3D clustering algorithms to our pre-processed TCGA and COSMIC dataset, namely HotSpot3D^9^, 3D hotspot^11^, HOTMAPS^13^, and PSCAN^10^. Given that these approaches compiled protein structures from different resources but they all used PDB, we restricted the focus to the 3D clusters identified based on PDB structures to make fair comparisons. For all four algorithms, default parameters were used if not specified. We applied HotSpot3D to our mutation data through the HotSpot3D web server^109^. For PSCAN, we tested both the mean and variance of the genetic effects within each scan window using the PSCAN R package, ultimately plotting the performance curve using the variance test results because they yielded a much better performance curve than the mean test. PSCAN input files were generated from SCORE-Seq^110^ as suggested. For the mutations in each gene, SCORE-Seq was applied to the corresponding genotypes in the affected tumor samples and their matched normal samples. When specifying SCORE-Seq parameters, we set the minor allele frequency (MAF) upper bound to 1 and minor allele count (MAC) lower bound to 0 to include all mutations. For HotMAPS, Tippett’s method was employed to aggregate the p-values of hotspot residues within each 3D cluster.

### Standard PPI network analyses

The initial amount of heat placed on each protein in G_PPI_ was determined by the number of tumor samples where the protein had mutation(s). Both in-frame and LOF mutations were accounted for. The weighting factors {*w*_*i,j*_} were all set to 1. The remaining settings were identical to thosevused in the heat diffusion model described earlier.

### Compiling catalytic residues

Catalytic residues were obtained from M-CSA^43^ (https://www.ebi.ac.uk/thornton-srv/m-csa/). We used the dataset that incorporates both the manually curated catalytic residues and their sequence homologs.

### Compiling benchmark gene sets

#### Known cancer genes

A list of 738 known cancer genes (tier 1 + tier 2) was obtained from CGC^49,50^ (https://cancer.sanger.ac.uk/census, 10/4/2023 release).

#### Non-cancer-associated genes

Non-cancer-associated genes were compiled from three sources: (i) 1,297 genes from Reva et al.^51^ in their category iv, i.e., genes with no functional mutations and no available associations with cancer; (ii) 129 genes annotated as “nonfunctional” by Saito et al.^52^, including genes frequently affected by passenger hotspot mutations and olfactory genes; 194 genes confidently under neutral selection in human cancers identified by Hess et al.^53^. By combining these three datasets we got a total of 1,574 unique genes, from which we removed 47 genes in CGC. The remaining 1,527 genes were considered non-cancer-associated.

### Compiling well-known cancer pathways and GO biological processes

#### Cancer signaling pathways

32 manually curated cancer signaling pathways were obtained from NetSlim^59^ (http://www.netpath.org/netslim). Specifically, we extracted DataNode from the GPML file of each pathway, from which we excluded DataNode without type or whose type is “Metabolite” or “Complex”.

#### General biological processes

7,530 Gene Ontology (GO) biological processes were obtained from the Molecular Signatures Database (MSigDB) C5 collection^54–58^ (http://www.gsea-msigdb.org/gsea/msigdb). We excluded GO biological processes that did not contain any protein-coding genes.

### Consistency with established biological processes

For every significantly interconnected module identified by NetFlow3D, we assessed the overlap between the module’s proteins and the genes of each GO BP, computing a Jaccard similarity coefficient. The alignment of a NetFlow3D-identified significantly interconnected module with established biological processes is determined by the highest Jaccard similarity coefficient between this module and any GO BPs. This criterion was also employed for evaluating randomly selected connected components and random groups of protein with significant 3D clusters.

### Mutation pattern analysis

Mutation enrichment was determined by the ratio of observed fraction of mutations over the relative length of proteins within each NetFlow3D module, well-known cancer pathway or GO BP. Relative length is defined as the sum of protein sequence lengths within each set divided by the total sequence length of all proteins with in-frame mutations.

### Definition of NetFlow3D-identified potential driver mutations

Within the significantly interconnected modules identified by NetFlow3D, we identified potential driver mutations: For each protein, we identified its most significant 3D cluster that surpasses the significance threshold and consider mutations within this cluster as potential driver mutations. This aligns with how we determine the initial heat score at each node. For each PPI, we identified the most significant 3D cluster that surpasses the significance threshold at its binding interface. The mutations within this cluster are designated as potential driver mutations. This aligns with how we determine heat propagation weight along each edge.

### Survival analysis

Patient clinical data were obtained from the TCGA Pan-Cancer Clinical Data Resource^111^ (TCGA-CDR). Patients without valid tumor status were excluded from the analysis. The overall survival (OS) data was used as the clinical outcome endpoint. Our analysis focused on the patients with in-frame mutations in our preprocessed TCGA pan-cancer dataset. We compared the overall survival between patients grouped by whether their mutations were identified as potential driver mutations by NetFlow3D. Kaplan-Meier estimation was used to generate survival curves for both groups. Cox regression was used to evaluate the statistical association between the presence of NetFlow3D-identified mutations and OS, with tumor stage, age, sex, and tumor mutation burden (TMB) included as covariates. For brain lower grade glioma (LGG), tumor stage was excluded from the Cox regression analysis due to unavailable data. The regression coefficients of NetFlow3D-identified mutations indicate their impact on hazard, with their exponential values representing the hazard ratio (HR) and p-values indicating the significance of the association. Furthermore, to focus on the survival impacts of mutations in the genes not curated by CGC, we treated NetFlow3D-identified mutations in known cancer genes and those not in known cancer genes as two independent predictors within a single Cox regression model. Besides, we re-generated survival curves after excluding patients with NetFlow3D-identified mutations in known cancer genes.

### Cloning and Plasmid Construction

Wild-type PPP2R1A clones, sourced from the hORFeome v8.1 collection^112^, were used as the template for site-directed mutagenesis conducted by Eurofins Scientific. The PPP2R1A_c.772C>T (p.Arg258Cys) mutation was introduced following the procedures outlined in our Clone-seq^41^ pipeline. We employed the Gateway cloning technology to insert the PPP2R1A or its mutant form into the pHAGE-CMV-GAW-3xFlag-IRES-PURO vector, and PPP2R3A into the pHAGE-CMV-GAW-3xMyc-IRES-PURO vector for subsequent analyses.

### Affinity purification

HEK 293T cells were maintained in DMEM medium supplemented 10% Fetal Bovine Serum. 8ug of PPP2R1A or PPP2R1A_c.772C>T were transfected into the cells with 40ul of 1 mg/ml-1 PEI (Polysciences, 23966) and 1.2ml OptiMEM (Gibco, 31085-062). After 48hrs incubation, cells were washed three times in 10ml DPBS (VWR, 14190144), resuspended in 500ul of RIPA buffer (50mM Tris pH7.5, 150Mm NaCl, 5mM EDTA, 1.0% NP-40, 0.25% Sodium Deoxycholate) and incubated on the ice for 30 minutes. The whole lysate is subjected to 120 seconds of 40% amplitude sonication using a sonifier cell disruptor (BRANSON,500-220-180). Centrifugation was used for 15 minutes at 16,100g and 4°C to separate the extracts. 500ul of cell lysate per sample reaction was incubated with 15ul of EZ view Red Anti-FLAG M2 Affinity Gel (Sigma, F2426) at 4°C overnight using a nutator in order to facilitate co-immunoprecipitation. Following incubation, bound proteins were eluted in 200ul of elution solution (10mM Tris-Cl pH 8.0, 1% SDS) at 65°C for 15 minutes after being washed three times in cell RIPA buffer.

### Cell culture, co-immunoprecipitation and western blotting

HEK 293T cells were cultured in 10 cm plates until they reached 40-50% confluency. 4 ug of bait construct (PPP2R1A or PPP2R1A_c.772C>T), 4 ug of prey construct (PPP2R3A), 40 ul of 1 mg/ml-1 PEI (Polysciences, 23966), and 1.2 ml of OptiMEM (Gibco, 31085-062) were used to transfect the cells. After 48hrs incubation, cells were washed three times in 10ml DPBS (VWR, 14190144), resuspended in 500ul RIPA buffer (50mM Tris pH7.5, 150Mm NaCl, 5mM EDTA, 1.0% NP-40, 0.25% Sodium Deoxycholate) and incubated on the ice for 30 minutes. Whole lysate is sonicated on a sonifier cell disruptor (BRANSON,500-220-180) for 120 seconds at 40% amplitude. Extracts were cleared by centrifugation for 15 minutes at 16,100g at 4°C. 500ul of cell lysate per sample reaction was incubated with 15ul of EZ view Red Anti-FLAG M2 Affinity Gel (Sigma, F2426) at 4°C overnight using a nutator. After incubation, bound proteins were eluted in 200ul of elution solution (10mM Tris-Cl pH 8.0, 1% SDS) at 65°C for 15 minutes after being washed three times in cell RIPA buffer. Following an 8% SDS-PAGE gel run on FLAG-co-purified samples, the proteins were transferred to PVDF membranes. Anti-FLAG (Sigma, F1804), and Anti-MYC (Invitrogen, 132500) at both 1:5000 dilutions were used for immunoblotting analysis.

### Proteomic sample preparation

IP eluates were subjected to reduction with 200 mM TCEP for 1h at 55°C. Subsequently, alkylation was performed for 30 minutes at room temperature in darkness using 375 mM iodoacetamide. The samples were then digested using Trypsin Gold, mass spectrometry grade (catalog no. V5280; Promega), at an enzyme-to-substrate ratio of 1:100. The samples were incubated overnight at 37°C. Following this, the concentrations of peptides were quantified using the Pierce Quantitative Colorimetric Peptide Assay (catalog no. 23275; Thermo Scientific). For TMT tests, samples were resuspended and normalized using 1M triethylammonium bicarbonate (catalog no. 90114; Thermo Scientific). Samples were labeled using TMT10plex Isobaric Mass Tagging Kit (catalog no. 90113; Thermo Scientific) at a (w/w) label-to-peptide ratio of 20:1 for 1h at room temperature. Labeling reactions were quenched by the addition of 5% hydroxylamine for 15 minutes and pooled and dried using a SpeedVac. Labeled peptides were enriched and fractionated using Pierce High pH Reversed-Phase Peptide Fractionation Kit according to the manufacturer’s protocol (catalog no. 84868; Thermo Scientific). Liquid chromatography–tandem mass spectrometry Fractions were analyzed using an EASY-nLC 1200 System (catalog no. LC140; Thermo Scientific) equipped with an in-house 3 μm C18 resin-(Michrom BioResources) packed capillary column (125 μm × 25 cm) coupled to an Orbitrap Fusion Lumos Tribrid Mass Spectrometer (catalog no. IQLAAEGAAPFADBMBHQ; Thermo Scientific). The mobile phase and elution gradient used for peptide separation were as follows: 0.1% formic acid in water as buffer A and 0.1% formic acid in 80% acetonitrile as buffer B; 0–5 min, 5%-8% B; 5–65 min, 8–45% B; 65–66 min, 45%-95% B; 66–80 min, 95% B; with a flow rate set to 300 nl min−1. MS1 precursors were detected at m/z = 375–1500 and resolution = 120,000. A CID-MS2-HCD-MS3 method was used for MSn data acquisition. Precursor ions with charge of 2+ to 7+ were selected for MS2 analysis at resolution = 50,000, isolation width = 0.7 m/z, maximum injection time = 50 ms and CID collision energy at 35%. 6 SPS precursors were selected for MS3 analysis and ions were fragmented using HCD collision energy at 65%. Spectra were recorded using Thermo Xcalibur Software v.4.4 (catalog no. OPTON-30965; Thermo Scientific) and Tune application v.3.4 (Thermo Scientific). Raw data were searched using Proteome Discoverer Software 2.3 (Thermo Scientific) against an UniProtKB human database.

### Downstream proteomic analysis

We employed our computational tool Magma^113^ to analyze mass spectrometry proteomics data. Magma quantifies the differences in protein abundance between two experimental conditions by calculating fold-change (FC) and p-values. By comparing each bait protein (PPP2R1A or PPP2R1A_c.772C>T) against untransfected HEK293T cells, we identified the bait protein’s interactors using criteria of fold change (FC) > 2, adjusted p-value < 0.05, and peptide-spectrum matches (PSM) > 10. Our analysis was then narrowed to the combined set of interactors for both PPP2R1A and its mutant form PPP2R1A_c.772C>T. To elucidate the specific effects of the c.772C>T mutation, we generated a volcano plot using the FC and adjusted p-values derived from the comparison between the mutant variant PPP2R1A_c.772C>T and the wild-type PPP2R1A. Known contaminants in AP-MS experiments, including keratin (KRT), myosins (MYO), small ribosomal subunit proteins (RPS), heat shock-related 70 kDa proteins (HSPA), and large ribosomal subunit proteins (RPL), were excluded from the analysis.

## Acknowledgments

The results shown here are based upon data generated by the TCGA Research Network: https://www.cancer.gov/tcga. We thank members of the Yu laboratory for helpful discussions and guidance; and Zui Tao for her suggestions on the conceptualization.

## Author contributions

Conceptualization: YZ, HY

Methodology: YZ, JB, AKL

Visualization: YZ, AKL, TQ, YS, JS

Formal analysis: YZ, AKL, LL, YS, JS, SG

Validation: YZ, JJK, GW, LC

Data curation: YZ, AKL, JZ, SW

Funding acquisition: HY Supervision: HY

Writing – original draft: YZ

Writing – review & editing: YZ, HY, AKL, LL, JJK, JZ, SW, YS

## Ethics declarations

### Data availability

The TCGA MC3 dataset was downloaded from https://gdc.cancer.gov/about-data/publications/mc3-2017. TCGA RNA-seq data and proteome profiling data was downloaded from https://portal.gdc.cancer.gov/. The ID mapping file was downloaded from https://ftp.uniprot.org/pub/databases/uniprot/current_release/knowledgebase/idmapping/by_organism/HUMAN_9606_idmapping.dat.gz. SIFTS data was downloaded from https://www.ebi.ac.uk/pdbe/docs/sifts/index.html. Protein Mass Spectrometry Raw Data is publicly available at MassIVE (https://massive.ucsd.edu/) under accession code MSV000094298. Source data and supplementary tables are available on Zenodo (https://doi.org/10.5281/zenodo.7680146).

### Code availability

A GitHub repository containing the source code of NetFlow3D is available (https://github.com/haiyuan-yu-lab/NetFlow3D).

