## Supplementary Notes and Figures for "A multiscale functional map of somatic mutations in cancer integrating protein structure and network topology"

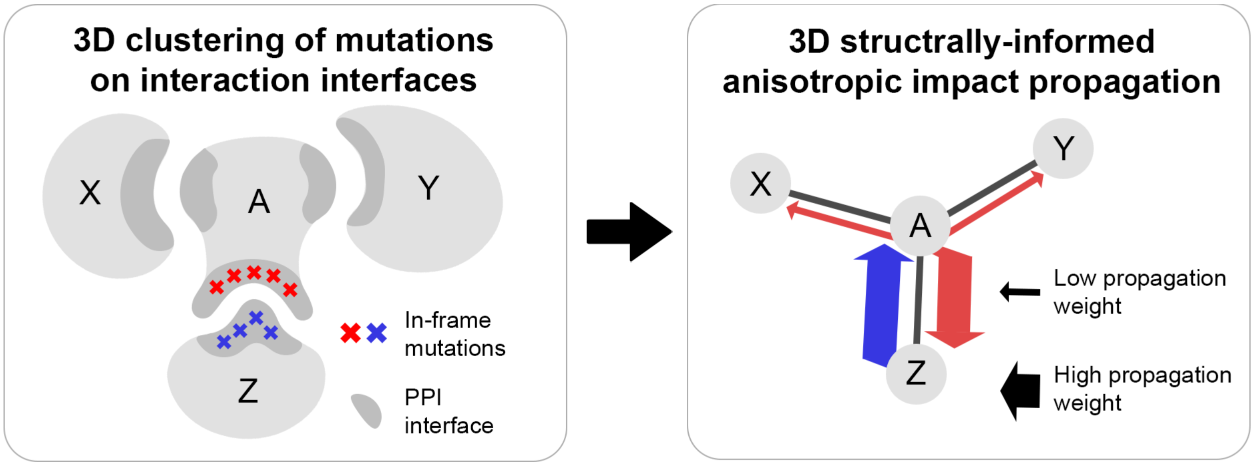


**Supplementary Fig. 1. NetFlow3D's weighted propagation strategy.** NetFlow3D recognizes that proteins often interact with different partners using distinct 3D structural interfaces. It, therefore, uses this information to weight the impact of spatially clustered mutations at a specific PPI interface on different interaction partners differently (anisotropic).

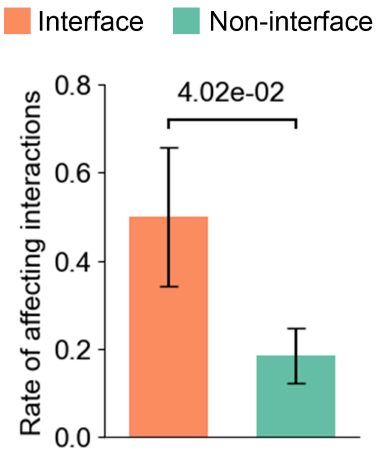


**Supplementary Fig. 2. The edgetic effect of somatic cancer driver mutations.** To validate the "edgetic effect" of functional somatic in-frame mutations, we experimentally generated 37 mutations in the significant 3D clusters identified by NetFlow3D, and tested their effects on protein-protein interactions via yeast two-hybrid (Y2H) assay (Supplementary Note). In total, we screened 48 mutation-interaction pairs, including 10 pairs where mutations were at the interaction interfaces and 38 pairs where mutations were not. The assay revealed perturbations in interactions for 5 pairs with interface mutations and 7 pairs with non-interface mutations, demonstrating that likely driver somatic mutations at interaction interfaces have a significantly higher rate of interaction-perturbing effects compared to non-interface mutations, thereby validating their "edgetic effect".

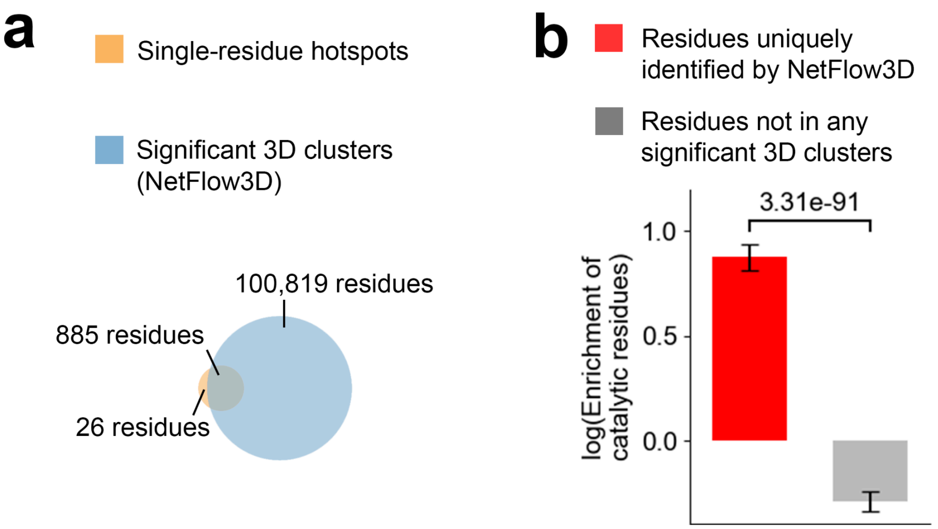


**Supplementary Fig. 3. Comparison of 3D clustering results from NetFlow3D with single-residue hotspots detected by the recurrence-based methods.** (**a**) Overlap between residues within significant 3D clusters identified by NetFlow3D and those identified as hotspots by the single-residue-based method. For visual clarity, the area proportions in the Venn diagram are scaled based on the square root of the residue count. (**b**) Enrichment analysis of catalytic residues focusing on the residues uniquely identified by NetFlow3D.

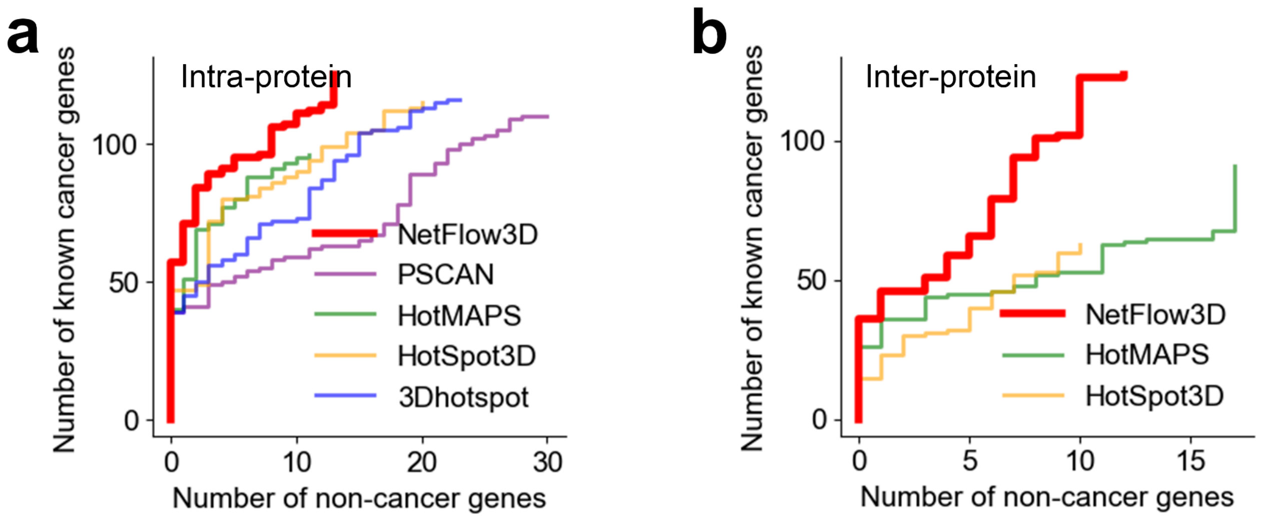


**Supplementary Fig. 4. Performance comparison between NetFlow3D and state-of-the-art 3D clustering algorithms on a COSMIC pan-cancer dataset completely independent from TCGA.** (**a**) Performance curves derived from intra-protein 3D clustering results. (**b**) Performance curves derived from inter-protein 3D clustering results.

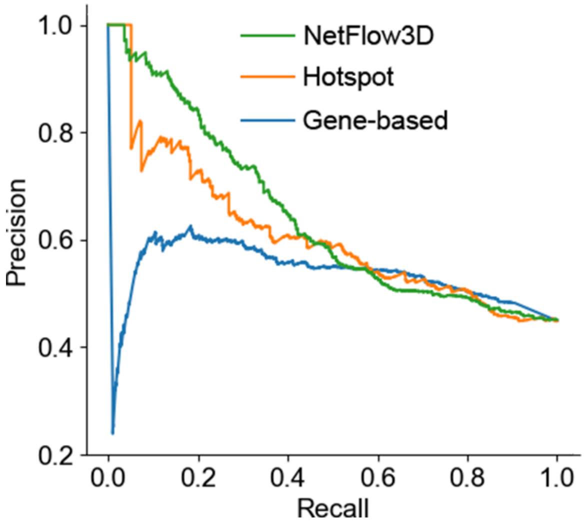


**Supplementary Fig. 5. Precision-recall (PR) curves of three methods for identifying cancer driver mutations.** Three methods focus on different test units: (i) 3D cluster (NetFlow3D), (ii) single residue (Hotspot), and (iii) whole gene (Gene-based). Gene-level precision-recall curves were generated using known cancer genes from the Cancer Gene Census (CGC) as positive labels and known non-cancer genes curated from the literature as negative labels. Genes with unknown labels were excluded to prevent potential misclassification. For the Hotspot and NetFlow3D methods, the most significant residue or 3D cluster was selected as the representative for each gene.

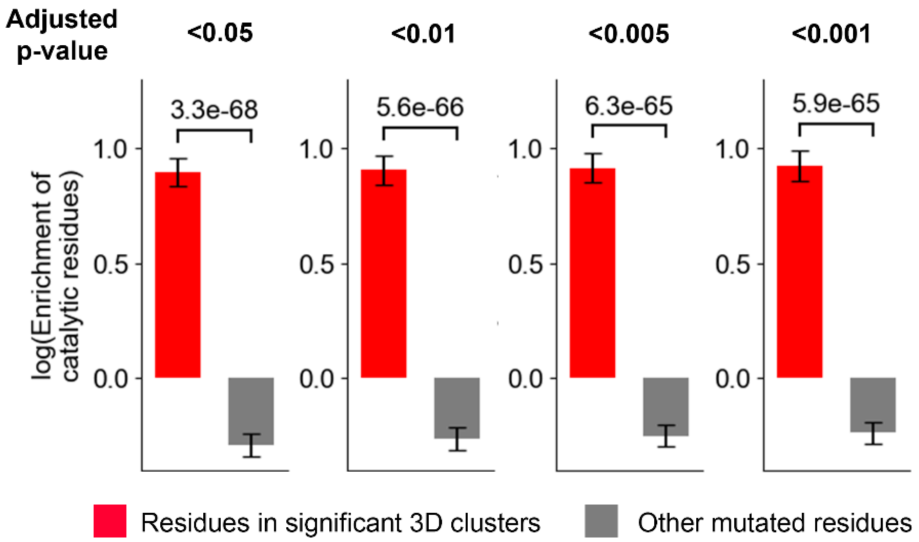


**Supplementary Fig. 6. Enrichment analysis of catalytic residues in significant 3D clusters across a range of statistical thresholds.** This analysis contrasts two residue groups: those within significant 3D clusters and other mutated residues not in significant 3D clusters. The p-values for comparisons were from Fisher's exact test.

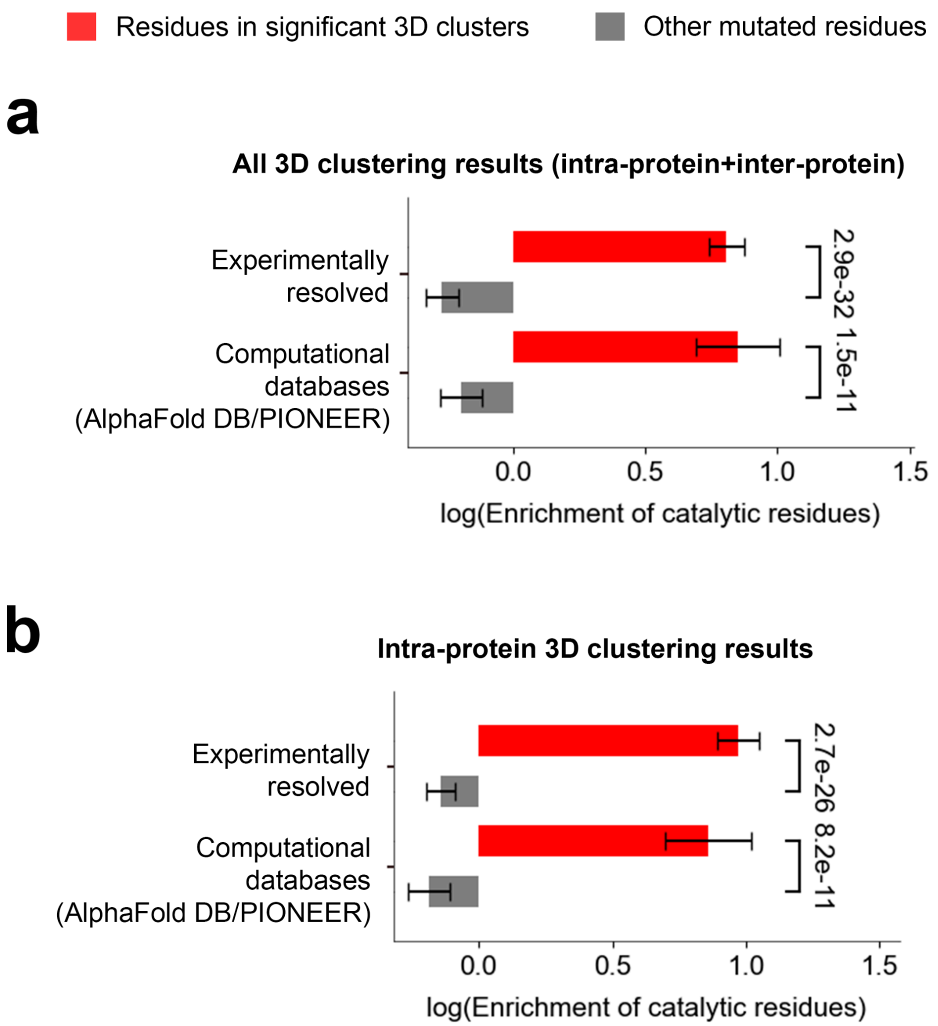


**Supplementary Fig. 7. Analysis of catalytic residue enrichment conducted distinctly on 3D clustering results derived from experimentally determined structures and deep-learning-generated 3D structural data.** (**a**) Utilizes results from both intra-protein and inter-protein 3D clustering analyses. (**b**) Focuses solely on the findings from intra-protein 3D clustering analysis.

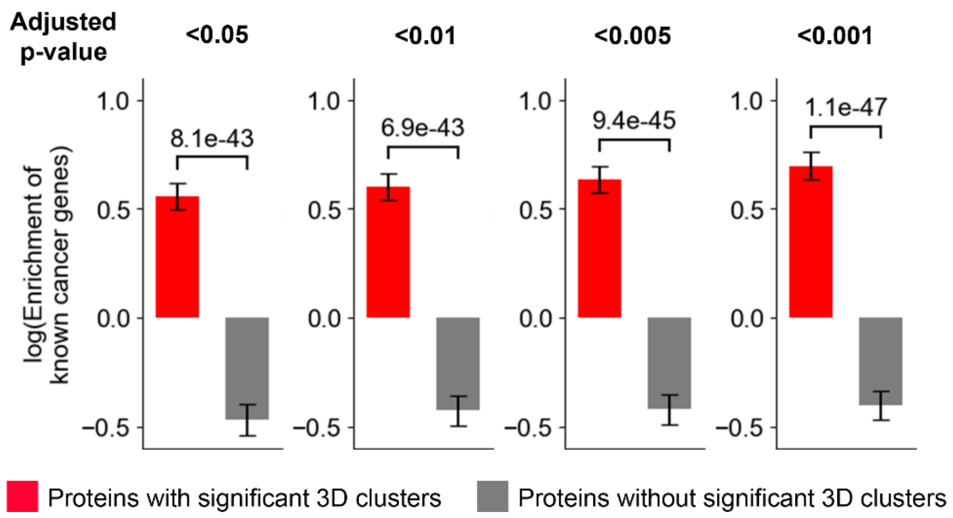


**Supplementary Fig. 8. Enrichment analysis of known cancer genes across two categories of proteins grouped by the presence of significant 3D clusters, with a range of statistical thresholds being applied.** The p-values for comparisons between proteins with and without significant 3D clusters were from Fisher's exact test.

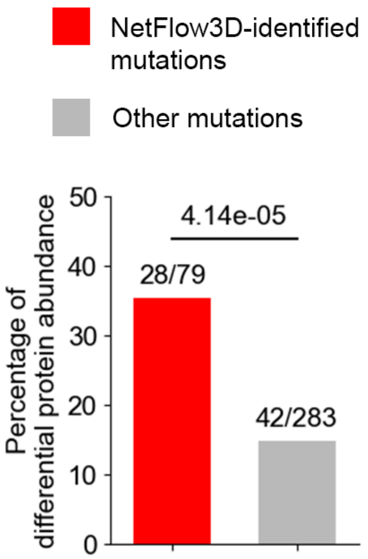


**Supplementary Fig. 9. Differential protein abundance analysis per gene and cancer type in scenarios with and without mutations of interest.** For each gene in each cancer type, the analysis required both scenarios to have at least 10 samples with available protein expression quantification data. The analysis was conducted for NetFlow3D-identified mutations (red) and other mutations (gray). The p-value was calculated using a two-proportion z-test.

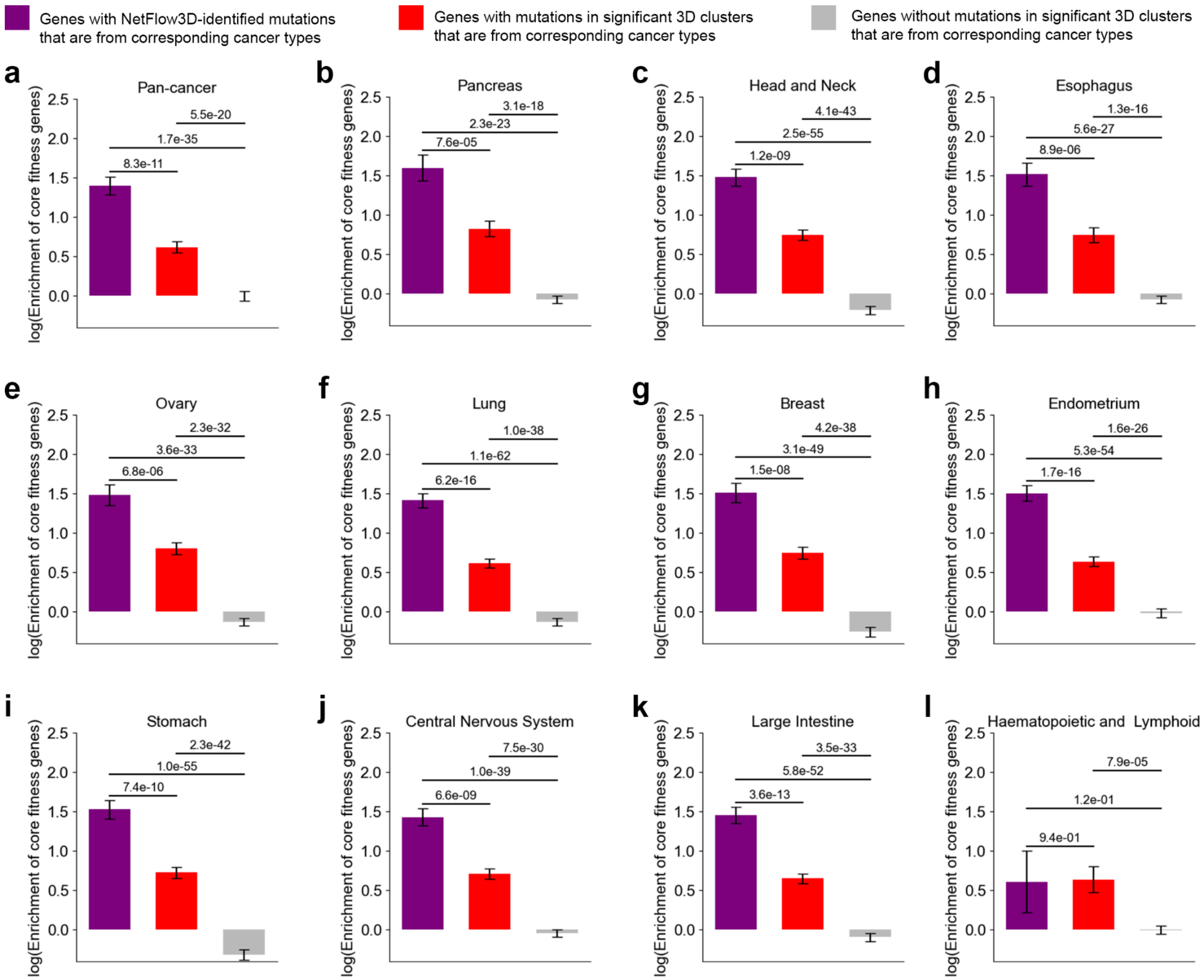


**Supplementary Fig. 10. Analysis of core fitness gene enrichment in NetFlow3D results.** Pan-cancer and cancer-type-specific core fitness genes were identified by Behan et al. using genome-scale CRISPR-Cas9 screening data in human cancer cell lines. The p-values were calculated using two-proportion z-tests.

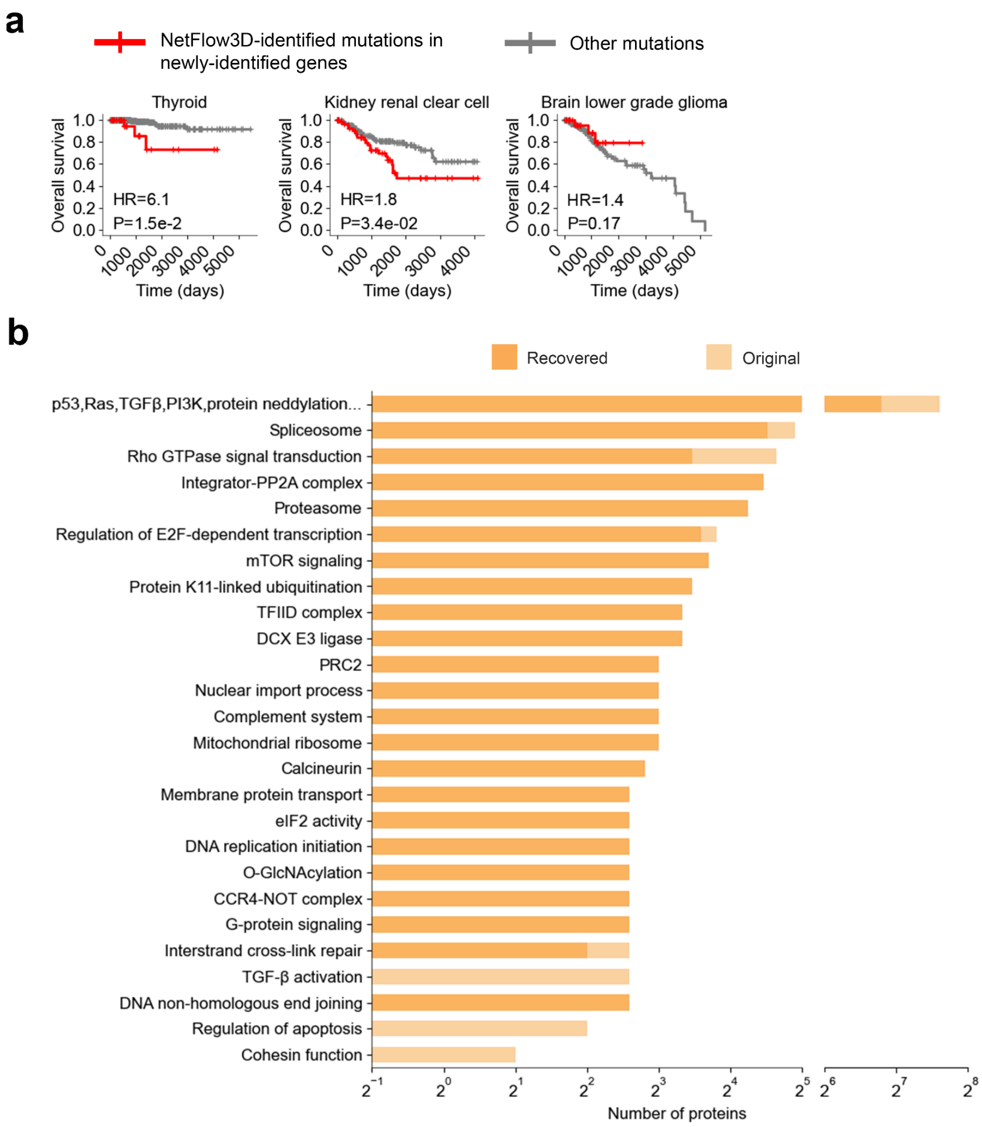


**Supplementary Fig. 11. Evaluation of NetFlow3D findings beyond known cancer genes.** (**a**) As an extension of Fig. 4d, the distinct survival impacts of NetFlow3D-identified mutations in genes not curated by CGC were evaluated in the same cancer types. Note that Adrenocortical carcinoma (ACC) was excluded from this analysis because it had fewer than 10 patients with NetFlow3D-identified mutations after excluding those with mutations in known cancer genes. (**b**) Mutation signals from known cancer genes were completely removed before re-applying the network propagation model (Supplementary Note). The x axis indicates the number of proteins within each original NetFlow3D module (light orange) and those that were recovered by the newly-identified significantly interconnected modules (dark orange). For a fair comparison, this recovery analysis focused exclusively on the proteins in the original NetFlow3D modules that are not known cancer genes.

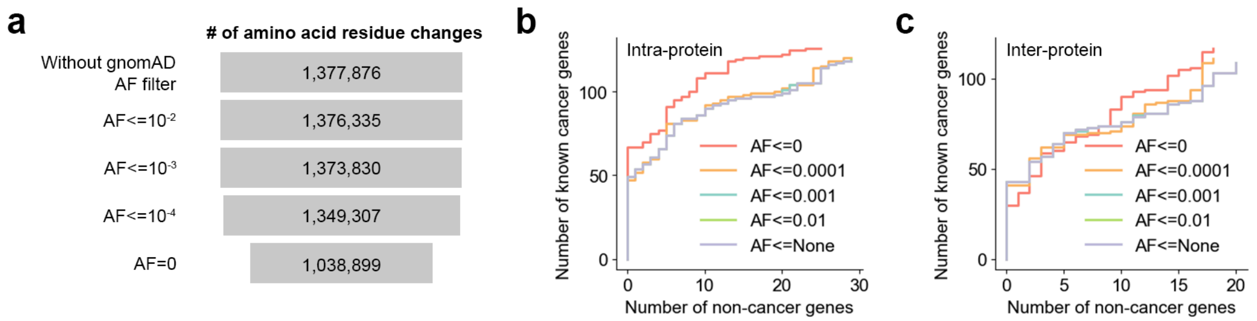


**Supplementary Fig. 12. Evaluating the outcomes of applying a range of germline variant filters at different gnomAD allele frequency thresholds.** (**a**) Number of preprocessed mutations using each germline variant filter. (**b**) Performance curves derived from intra-protein 3D clustering results. (**c**) Performance curves derived from inter-protein 3D clustering results.

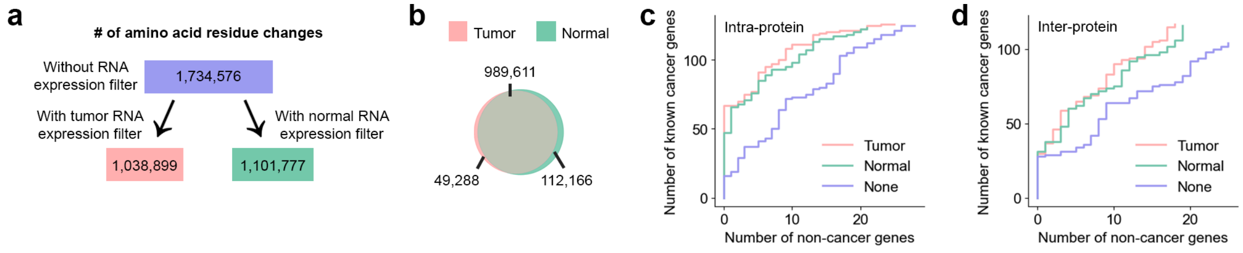


**Supplementary Fig. 13. Consequences of three RNA expression filtering approaches.** We evaluated the consequences of applying a tumor RNA expression filter (used by NetFlow3D), a normal RNA expression filter, and the scenario where no RNA expression filter is applied. For the normal RNA expression filter approach, we only retained the mutations in those genes with RNA expression levels >=1 FPKM in >=80% of normal tissue samples near the tumors of the corresponding cancer types. (a) Number of preprocessed mutations under each filtering scenario. (b) A high degree of overlap in the mutations that were retained after applying the tumor and normal RNA expression filter. (c-d) Analysis of these mutations using NetFlow3D revealed that both the tumor and normal RNA expression filters significantly improved performance.

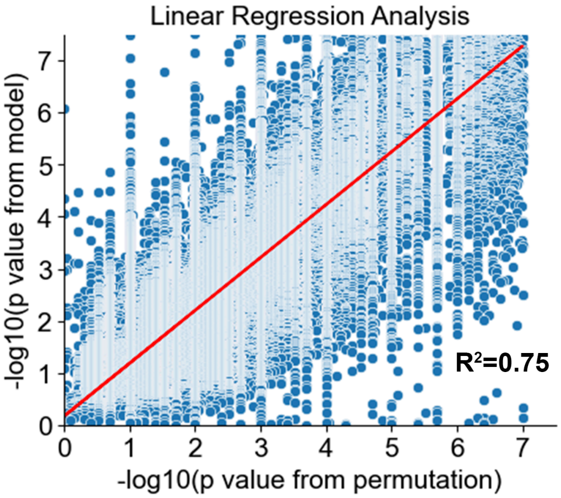


**Supplementary Fig. 14. Correlation analysis of 3D cluster p-values derived from permutation tests and NetFlow3D.** Each dot corresponds to a 3D cluster.

**Supplementary Note**

Linear regression analysis of protein abundance

TCGA proteome profiling data was obtained from Repository in the GDC data portal[1] (https://portal.gdc.cancer.gov/). Protein abundance for each gene in a sample was defined as the maximum quantification value among all its protein peptides. We used a linear regression model to evaluate the statistical association between NetFlow3D-identified mutations and protein abundance. This model controlled for gene-specific and tissue-specific baseline expression levels, as well as clinical covariates including sex, age, tumor stage, and TMB (Model: protein abundance ~ C(NetFlow3D-identified mutations) + C(gene ID) + C(cancer type) + age + C(sex) + TMB + C(tumor stage)). As a negative benchmark, we repeated the analysis by replacing C(NetFlow3D-identified mutations) with C(other mutations).

Re-applying network propagation model after excluding the mutation signals from known cancer genes

3D clusters composed exclusively of mutations in known cancer genes, as well as LOF enrichment signals within known cancer genes, were excluded from the analysis. Subsequently, initial heat scores and propagation weights were re-calculated following equation (13-16) (Methods), and the network propagation model was re-applied. We then compared the newly-identified significantly interconnected modules to the original NetFlow3D modules to assess the recovery rate of the initial NetFlow3D modules.

Proteoform diversity

To evaluate the impact of incorporating proteoform diversity, we applied our revised NetFlow3D framework to the same TCGA mutation dataset, this time mapping mutations to all possible isoforms. This thorough mapping approach yielded 3.5 times more amino acid residue changes (2.95M) than the canonical mapping approach (0.85M). We conducted 3D clustering analysis using AlphaFold DB 3D structures, which encompass ~186k human protein isoforms. This analysis was not applicable to PDB or PIONEER, give that 96% (7,195 out of 7,515) of human UniProt entries covered by PDB are canonical isoforms, and 95% (113,949 out of 119,526) of high-quality binary human PPIs in HINT are between canonical isoforms. The 3D clustering results are provided in Supplementary Table 11-12. Our analysis revealed that for 84% of genes (12,167 out of 14,455), the most significant 3D cluster was still identified from their canonical isoforms.

Permutation tests

To benchmark the p-values of 3D clusters in NetFlow3D via permutation tests, we conducted 107 permutations. In each permutation, we randomly shuffled the mutations per patient per protein, and then counted how many patients had mutations within each 3D cluster. These permutations gave us empirical distributions and p-values. Consequently, there is a strong correlation between the p-values from permutation tests and those from NetFlow3D, with R2=0.75 (Supplementary Fig. 11).

Mutagenesis experiments

We performed mutagenesis experiments where we introduced 38 NetFlow3D-identified potential driver mutations into the proteins according to our Clone-seq pipeline[2], and tested their effects on protein-protein interactions via our high-throughput yeast two-hybrid (Y2H) assay. In total, we screened 48 mutation-interaction pairs, including 10 pairs where mutations were at the interaction interfaces and 38 pairs where mutations were not.

Y2H assay

Y2H was performed as previously described[3]. Gateway LR reactions were used to transfer all wild-type/mutant clones into our Y2H pDEST-AD and pDEST-DB vectors. All DB-X and AD-Y plasmids were transformed into the Y2H strains MATα Y8930 and MATa Y8800, respectively. Thereafter, each of the DB-X MATα transformants (wild-type and mutants) were mated with corresponding AD-Y MATa transformants (wild-type and mutants) individually through automated 96-well procedures, including inoculation of AD-Y and DB-X yeast cultures, mating on YEPD media (incubated overnight at 30°C), and replica-plating onto selective Synthetic Complete media lacking histidine, leucine, and tryptophan, and supplemented with 1 mM of 3-amino-1,2,4-triazole (SC-Leu-Trp-His+3AT), SC-Leu-His+3AT plates containing 1 mg/l cycloheximide (SC-Leu-His+3AT+CHX), SC-Leu-Trp-Adenine (Ade) plates, and SC-Leu-Ade+CHX plates to test for CHX-sensitive expression of the LYS2::GAL1-HIS3 and GAL2-ADE2 reporter genes. The plates containing cycloheximide were used to select for cells that do not have the AD plasmid due to plasmid shuffling. Spontaneous auto-activators[4], therefore, were identified by growth on these control plates. These plates were incubated overnight at 30 °C and "replica-cleaned" the following day. Subsequently, plates were incubated for three more days, after which positive colonies were scored as those that grow on SC-Leu-Trp-His+3AT and/or on SC-Leu-Trp-Ade, but not on SC-Leu-His+3AT+CHX or on SC-Leu-Ade+CHX. Disruption of an interaction by a mutation was defined as at least 50% reduction of growth consistently across both reporter genes when compared to Y2H phenotypes of the corresponding wild-type allele as benchmarked by 2-fold serial dilution experiments. All Y2H experiments were repeated 3 times.
